# Disruption of putrescine export in experimentally evolved *Ralstonia pseudosolanacearum* enhances symbiosis with *Mimosa pudica*

**DOI:** 10.1101/2025.04.16.649159

**Authors:** Anne-Claire Cazalé, Marvin Navarro, Ginaini Grazielli Doin de Moura, David Hoarau, Floriant Bellvert, Sophie Valière, Caroline Baroukh, Philippe Remigi, Alice Guidot, Delphine Capela

## Abstract

Polyamines are essential molecules across all domains of life, but their role as signaling molecules in host-microbe interactions is increasingly recognized. However, because they are produced by both the host and the microbe, their dual origin makes their functional dissection challenging. The plant pathogen *Ralstonia pseudosolanacearum* GMI1000 secretes large amounts of putrescine both *in vitro* and in the xylem sap of host plants. In this study, we investigated the genetic changes underlying its experimental evolution into a legume symbiont. We showed that the *paeA* gene (RSc2277), which was repeatedly mutated during this process, encodes a putrescine exporter. Mutations in *paeA* completely abolished putrescine excretion *in vitro* and enhanced bacterial proliferation within nodules during interaction with the legume *Mimosa pudica*. When these mutations occurred in symbionts already capable of intracellular infection, it further increased bacterial load in nodules and allowed the detection of nitrogenase activity. In addition, *paeA*-mutated symbionts modulated host gene expression towards a more functional symbiotic state by repressing defense-related genes and inducing nodule development genes. These nodule development genes include genes encoding leghemoglobins and an arginine decarboxylase, a key enzyme in plant putrescine biosynthesis. These results indicate that bacterial and plant putrescine have distinct functions in legume symbiosis and highlight the complex role of polyamines in plant-microbe interactions.

**Importance:** Rhizobia, the nitrogen-fixing symbionts of legumes, emerged through repeated and independent horizontal transfers of some essential symbiotic genes. However, these transfers alone are often insufficient to convert the recipient bacterium into a functional legume symbiont. In a laboratory experiment, we evolved the plant pathogen *Ralstonia pseudosolanacearum* into a nodulating and intracellularly infecting symbiont of *Mimosa pudica*. This transition required genomic modifications in the recipient bacterium to activate its acquired symbiotic potential. Here, we demonstrated that one of these key adaptive modifications is the inactivation of bacterial putrescine export. This polyamine, when produced by the microsymbiont, appears to act as a negative signal for the plant. This study provides new insights into the distinct roles of bacterial- and plant-derived putrescine in plant-microbe interactions, highlighting their functional divergence despite being produced by both organisms.

## INTRODUCTION

Bacteria known as rhizobia establish mutualistic interactions with legumes, resulting in the formation of specialized plant organs called nodules. Within these nodules, hundreds of thousands of bacteria reside intracellularly, fixing atmospheric nitrogen into ammonia to benefit the plant. In return, bacteria receive carbon sources from the plant along with protection from the external environment. To initiate this interaction, most rhizobia synthesize lipochitooligosaccharides, called Nod Factors, which are specifically recognized by plant receptors and trigger nodule organogenesis and early infection processes (1–5). Bacteria penetrate the roots either through crack entry (intercellular infection) or by attaching to root hairs, which subsequently invaginate to form infection threads (6–8). These infection threads progress through the root cell layers, guiding the bacteria towards the developing nodule. Then, bacteria are released into the cytoplasm of nodule cells via an endocytosis-like process and differentiate into nitrogen-fixing bacteroids. Intracellular bacteroids are enclosed by a plant-derived membrane, forming structures known as symbiosomes (9, 10). Within symbiosomes, bacteroids face challenging conditions, including highly acidic pH, extremely low oxygen concentrations, the presence of reactive oxygen species and likely high osmotic pressure. All stages of this symbiotic interaction are tightly regulated by mutual recognition processes (11). While the bacterial genes involved in the early stages of symbiosis have been well studied (12, 13), those required for the later stages, such as the survival and persistence of bacteria within nodule cells, remain far less understood. This is particularly the case outside terminally differentiated systems specific to certain legumes, such as *Medicago* and *Pisum* in the IRLC (Inverted Repeat-Lacking Clade) or *Aeschynomene* species in the Dalbergioid clade (14, 15). In these interactions, the proliferation and differentiation of intracellular bacteria is controlled by plant peptides called Nodule-specific Cysteine-Rich (NCR) or NCR-like peptides (15, 16). These peptides interfere with various bacterial cellular processes, inhibiting cell division while promoting nitrogen fixation (17–21). In such cases, symbiotic bacteria must possess the specific transporters, BacA or BclA (22–24), or modified lipopolysaccharides (LPS) (25), or peptidoglycan-modifying enzymes (26) to survive within nodule cells.

Rhizobia are polyphyletic bacteria that belong to 21 different genera and hundreds of species among two classes of proteobacteria, alpha and beta (27). The symbiotic capacity of these bacteria emerged following independent and repeated horizontal transfers of symbiotic genes essential for the production of Nod factors (*nod* genes) and the synthesis and functioning of nitrogenase (*nif* and *fix* genes). However, the transfer of *nod, nif* and *fix* genes are not always sufficient to convert a strain that receives these genes into a functional nitrogen-fixing legume symbiont (28–31). In a previous evolution experiment, we transferred the symbiotic plasmid of *Cupriavidus taiwanensis* LMG19424, a natural symbiont of *Mimosa pudica*, into the plant pathogenic bacterium *Ralstonia pseudosolanacearum* GMI1000. The resulting chimeric strain was unable to nodulate *M. pudica.* However, after multiple large-scale inoculation trials involving hundreds of plants, three nodules appeared, from which we isolated three independent nodulating variants of *R. pseudosolanacearum* (32). These variants acquired the ability to nodulate *M. pudica* through mutations that inactivated the major determinant of *R. pseudosolanacearum* pathogenicity, its type III secretion system (T3SS). Mutations that conferred nodulation affected either the master regulator *hrpG* (33) or the T3SS structural gene *hrcV*. The three nodulating strains were then submitted to serial cycles of nodulation on *M. pudica*. The symbiotic properties of bacteria improved rapidly during the first cycles and then slowed down (34). After 35 evolution cycles, nodulation competitiveness was almost equivalent to that of the natural rhizobium *C. taiwanensis*, while bacterial proliferation within the nodules, although greatly improved over the cycles, did not reach the level of *C. taiwanensis* and nitrogen fixation was not achieved. Strong adaptive mutations improving the symbiotic properties of *R. pseudosolanacearum* were previously identified. In particular, intracellular infection was enhanced through mutations in the EfpR or PhcA regulatory pathways (32, 35, 36). EfpR and PhcA are master regulators controlling hundreds of genes, either positively or negatively (35, 37–41), with approximately 160 genes being commonly regulated by both. These shared targets include genes involved in EPS synthesis and motility, hemin/siderophore transport and metabolism genes, as well as genes encoding Hrp and T3 effector proteins. Additionally, mutations in *phcA* and *efpR* have been shown to broadly activate bacterial metabolic activities (35, 37, 38, 42). On the plant side, the analysis of *M. pudica* gene expression profiles in response to progressively adapted *R. pseudosolanacearum* strains revealed a correlation between bacterial adaptation and a gradual increase in the number of plant genes differentially expressed that are also differentially expressed during the interaction with the natural symbiont *C. taiwanensis* strain (43).

In this study, we continued the evolution experiment until 60 cycles in order to further enhance the symbiotic capacities of bacteria. Among the genes that were repeatedly mutated in this experiment, we identified the RSc2277 gene, which was mutated six times independently. Interestingly, low levels of nitrogenase activity could be detected in nodules formed by some evolved clones carrying a mutation in this gene. We showed that the inactivation of RSc2277 is highly adaptive, as it significantly increased bacterial proliferation within nodules and led to the detection of nitrogenase activity. The RSc2277 gene encodes a protein similar to the PaeA proteins of *Salmonella typhimurium* and *Escherichia coli.* In these bacteria, PaeA has been shown to function as a cadaverine/putrescine exporter, which is critical for bacterial survival under some stress conditions, likely by reducing toxic levels of intracellular polyamines and maintaining cation homeostasis (44, 45). *R. pseudosolanacearum* GMI1000 is known to synthesize and export large quantities of putrescine in its natural habitat the xylem of tomato (46, 47), as well as in cultures on various carbon sources (42, 48). However, the mechanisms of putrescine export in *R. pseudosolanacearum* have not been evidenced. Here, we show that inactivation of a PaeA homologue in *R. pseudosolanacearum* completely abolished putrescine export in culture and improved symbiosis with *M. pudica*.

## RESULTS

### The *paeA* (RSc2277) gene is repeatedly mutated in four independent parallel lineages of *R. pseudosolanacearum* experimentally evolved into legume symbionts

In a previous study we described the dynamics of occurrence of mutations in *R. pseudosolanacearum* populations of five lineages (B, F, G, K, M) evolved for 35 nodulation cycles on the legume *Mimosa pudica*. This analysis revealed genes that were mutated at higher frequency than expected by chance. Among the most frequently mutated genes, the gene RSc2277 was found mutated in four evolved clones from three lineages (lineages G, K and M). These four mutations were fixed in populations, which can be indicative of their adaptive character (Table 1). Indeed, in the G lineage, Doin de Moura et al. (34) previously showed that the V321G mutation in RSc2277, occurring in cycle 5, was highly adaptive, increasing bacterial *in planta* fitness by over 100-fold. When we continued the evolution of the five lineages up to 60 cycles of nodulation, two new mutations in RSc2277 occurred in evolved clones of the B lineage at cycles 41 and 45 (Fig. 1, Table 1). The *R. pseudosolanacearum* RSc2277 protein contains a signal peptide, three transmembrane domains, two tandem cystathionine beta-synthase (CBS) domains, and a CorC-HlyC C-terminal domain (Fig. S1C). The closest functionally characterized homolog of RSc2277 is the PaeA proteins from *Salmonella enterica* serovar Typhimurium and *E. coli*, which share 48% and 46% sequence identity with RSc2277, respectively, and display a similar 3-dimensional structure (Fig. S1AB). In *S.* Typhimurium and *E. coli*, PaeA proteins have been shown to function as polyamine export proteins (44, 45). In the following, we have named the RSc2277 gene *paeA*.

**Fig. 1.**
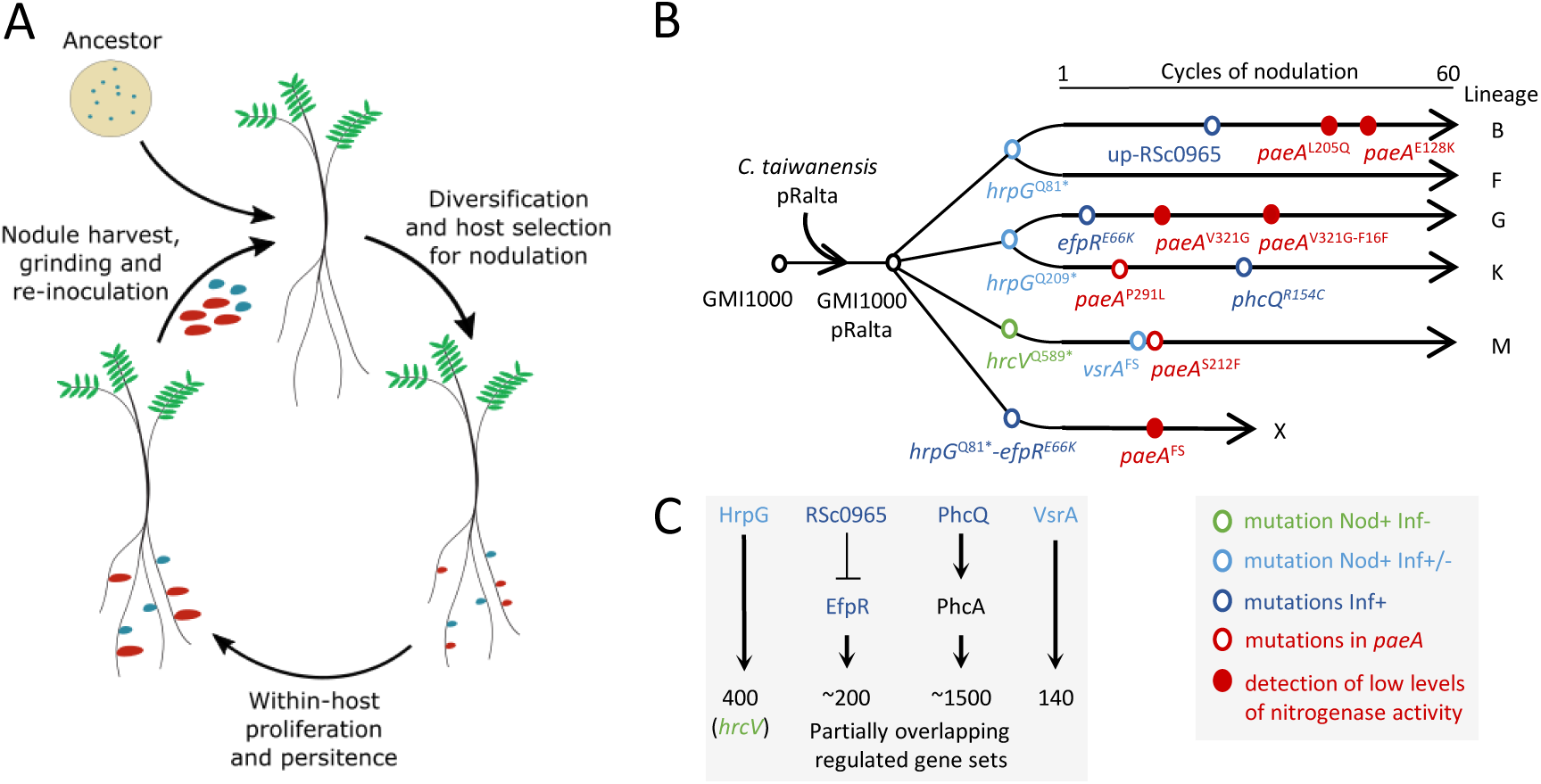
Occurrence of *paeA* mutations during the experimental evolution of *R. solanacearum* into *Mimosa pudica* symbionts. (**A**) Overview of the evolution experiment. In each cycle, inoculated bacteria diversify due to a transient hypermutagenesis phenomenon that occurs in the rhizosphere (50), and the most competitive variants for host entry are selected by the plants to form nodules (34). Within the nodules bacteria multiply before the nodules are harvested and ground to inoculate plants of the next cycle. (**B**) To initiate the evolution experiment, the symbiotic plasmid pRalta of *C. taiwanensis* was introduced into the GMI1000 strain of *R. pseudosolanacearum*. Three first nodulation variants were obtained when the resulting strain GMI1000 pRalta was massively inoculated onto *M. pudica* (32), then 5 independent lineages (B, F, G, K, M) were derived from these three ancestors and evolved for 60 cycles. The sixth lineage X was derived from the *hrpG*^Q81*^*-efpR*^E66K^ reconstructed mutant and evolved for 15 cycles. Red circles indicate the occurrence of *paeA* mutations along the experiment. In some cases, this mutation was associated with the detection of nitrogenase activity (filled red circles). The main adaptive mutations allowing nodulation (*hrpG*^Q81*^, *hrpG*^Q209*^ and *hrcV*^Q589*^ stop mutations) and intracellular infection (*hrpG*^Q81*^, *hrpG*^Q209*^, up-RSc0965, *efpR*^E66K^, *phcQ*^R154C^ and *vsrA*^FS^) were identified (32, 35, 36, 49). (**C**) A simplified schematic of *the R. pseudosolanacearum* regulatory pathways involving these mutations is provided. The HrpG, EfpR, PhcA and VsrA regulons have been identified and are partially overlapping (33, 35, 37–41). *, stop mutation. up, intergenic mutation upstream the start codon. FS, frameshift mutation.

**Table 1.**
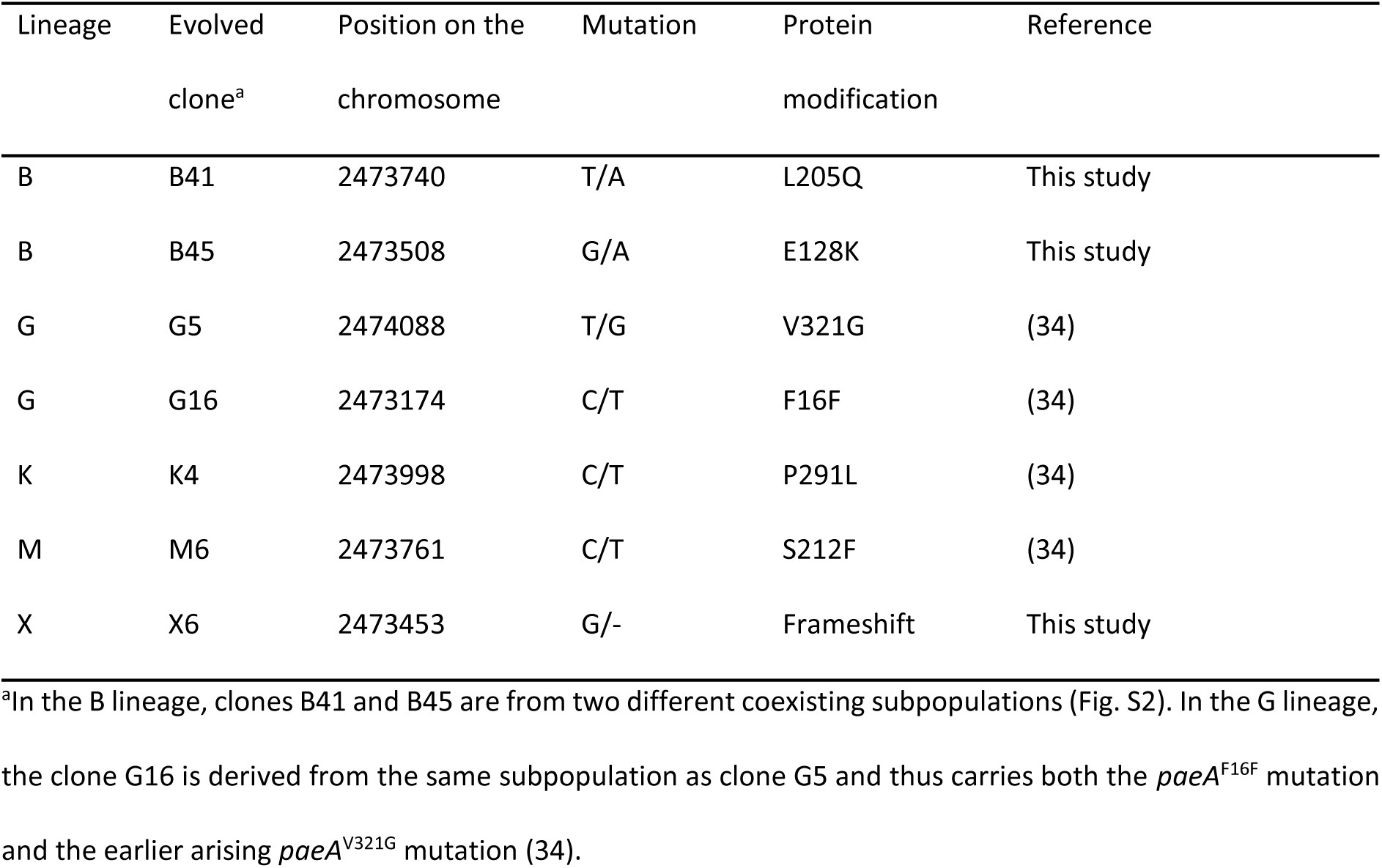
Mutations in the *paeA* gene occurred during the experimental evolution of *R. pseudosolanacearum*.

### Nitrogenase activity was detected in some *Rastonia* evolved clones mutated in *paeA*

Mutualism was not achieved in any of the five lineages even after 60 cycles of evolution on *Mimosa pudica*. However, low levels of nitrogenase activity measured by Acetylene Reduction Assays (ARA) were detected in nodules formed by some evolved clones. Interestingly, higher levels of nitrogenase activity were specifically detected in evolved clones B41, B45 and G5, all three carrying a mutation in *paeA*. However, no nitrogenase activity was detected in clones K4 and M6, both also carrying a mutation in *paeA*. One hypothesis is that the different mutations do not have the same consequence on PaeA protein function. However, the more appealing hypothesis was that *paeA* could trigger an increase in nitrogenase activity depending on the bacterial genetic background in which these mutations occurred. Indeed, in the nitrogen-fixing clones B41, B45 and G5, the *paeA* mutations occurred in evolved bacteria that infect nodule cells very well due to mutations in both *hrpG* (Q81* or Q209* stop mutations) and in the *efpR* pathway (mutations up-RSc0965 or *efpR*^E66K^) (35) (Fig. 1). While in the non-fixing evolved clones K4 and M6, mutations in *paeA* occurred in evolved bacteria that infect nodule cells only partially due to mutations in *hrpG* (Q81* or Q209* stop mutations) (32) or *hrcV* (Q589* stop mutation) and *vsrA* (frameshift) (49), respectively (Fig. 1). In the K lineage, a mutation improving intracellular infection (the mutation *phcQ*^R154C^) cumulated with the *paeA* mutation in the clone K13, but no nitrogenase activity was detected in nodules induced by this clone (Fig. 2). Moreover, we started a new evolution lineage (lineage X), this time using as ancestor a reconstructed mutant GMI1000 pRalta *hrpG*^Q81*^ *efpR*^E66K^ that infects well nodule cells. After a few cycles, a frameshift mutation in *paeA* occurred and was again associated with detectable levels of nitrogenase activity in nodules (clone X6, Fig. 2).

**Fig. 2.**
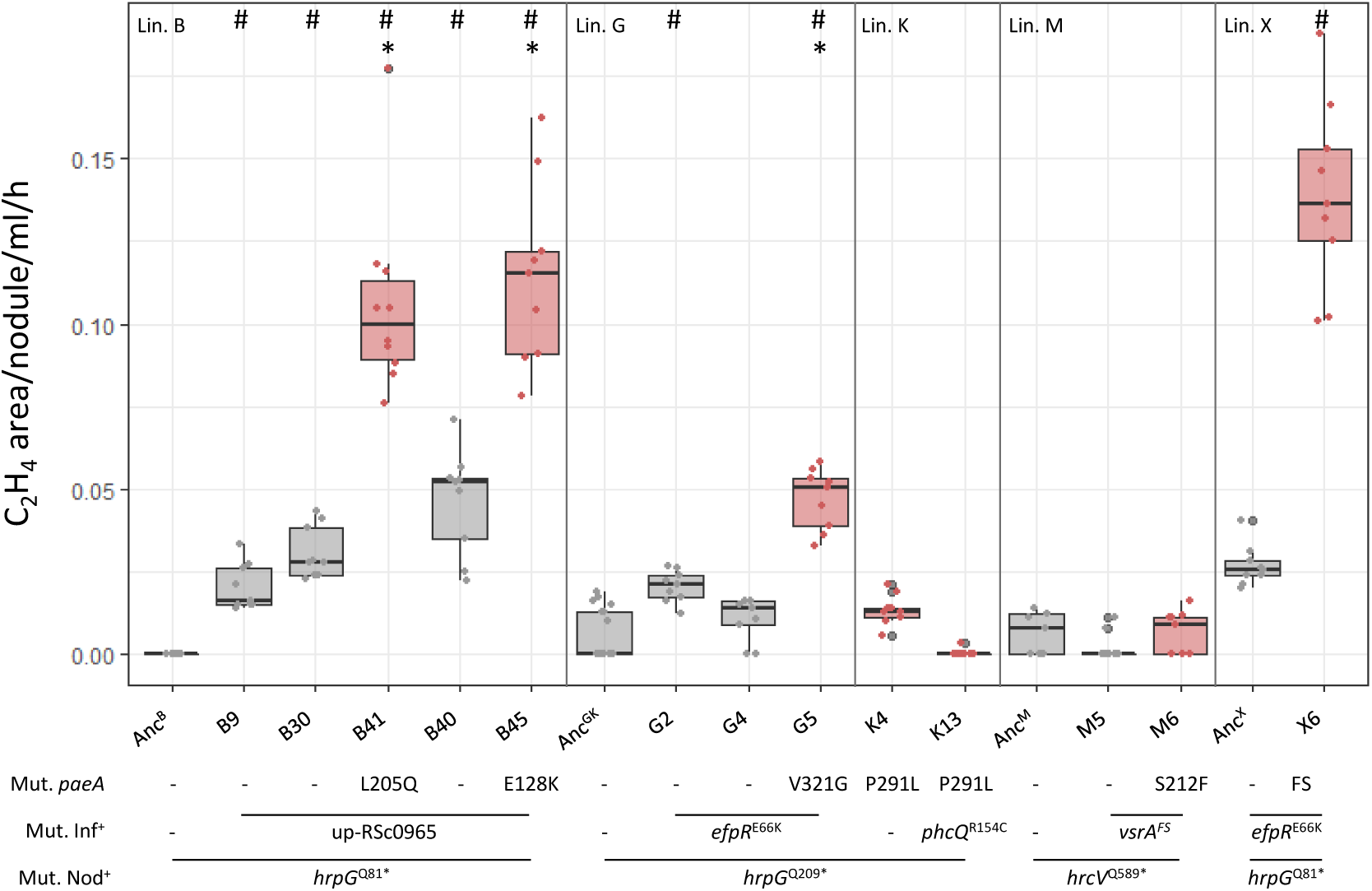
Nitrogenase activity in nodules formed by *Ralstonia* evolved clones. Acetylene reduction assays were performed on plants inoculated with isolated evolved clones 15 days after inoculation. Red box plots correspond to evolved clones carrying a mutation in *paeA*. Each measurement was taken from a pool of 6 plants. At least three measurements per experiment and three independent experiments were performed per strain. Lin., lineage. Anc^B^, Anc^GK^, Anc^M^, Anc^X^, nodulating ancestors of the B, G and K, M or X lineages. Mut. Nod^+^, mutation conferring the nodulation capacity. Mut. Inf^+^, main mutation conferring the capacity to infect nodules intracellularly. FS, frameshift mutation. The clone B30 is the closest evolved ancestor of clone B41 and the clone B40 is the closest evolved ancestor of clone B45 (see the phylogeny of evolved clones of the B lineage in Fig. S2). The complete list of mutations present in evolved clones is provided in Table S3. #Significantly different from the nodulating ancestor, *significantly different from the closest evolved ancestor (*P*<0.05, pairwise Wilcoxon test).

### Inactivation of *paeA* allows the detection of low levels of nitrogenase activity in *Ralstonia* clones mutated in the *efpR* or *phc* regulatory pathways

Evolved clones have accumulated many mutations during the evolution experiment (34, 50). In order to know whether the *paeA* mutation is responsible for the detected nitrogenase activity, we reconstructed the *paeA*^V321G^ mutation and an unmarked deletion of *paeA* in the strain GMI1000 pRalta carrying the *hrpG*^Q81*^ and *efpR*^E66K^ mutations, which confer nodulation and nodule intracellular infection capacity, respectively (32, 35). We measured the nitrogenase activity by ARA in nodules induced by these mutants. At 15 days post inoculation (dpi), we detected nitrogenase activity in both *paeA*^V321G^ and *paeA* deletion mutants but not in the parental strain (GMI1000 pRalta *hrpG*^Q81*^ *efpR*^E66K^). This shows that mutations in *paeA* in this background allowed the detection of nitrogenase activity in nodules and that the *paeA*^V321G^ mutation is equivalent to a loss of function (Fig. 3A). No nitrogenase activity was detected in the same strain in which we deleted the *nifH* gene, meaning that the ethylene measured in ARA is indeed due to the functioning of nitrogenase. Moreover, complementing the *paeA*^V321G^ mutant with the wild-type allele abolished nitrogenase activity (Fig. S3A).

**Fig. 3.**
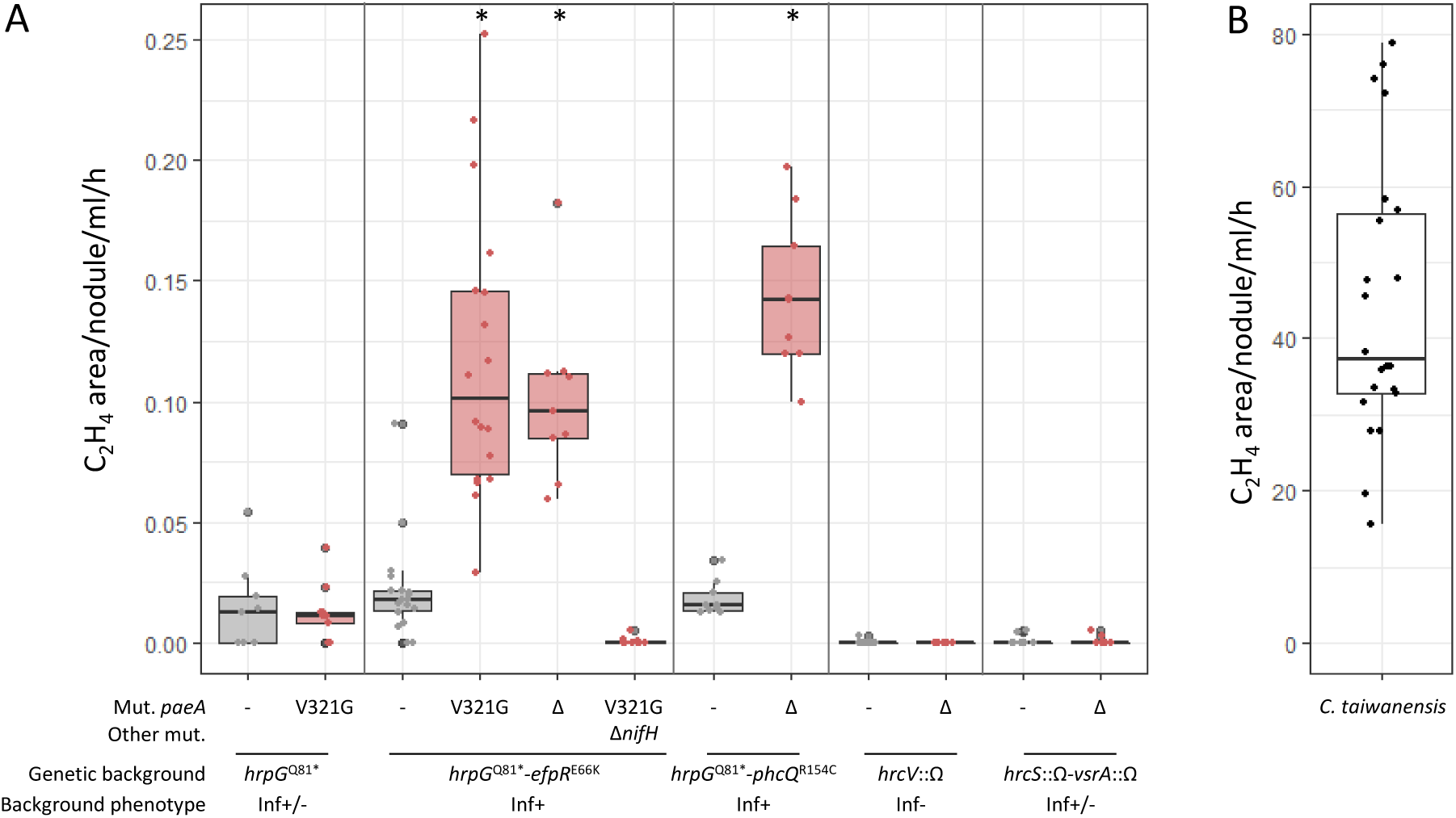
Acetylene reduction assays of plants inoculated with *Ralstonia paeA* reconstructed mutants (A) and *C. taiwanensis* (B) 15 days post inoculation. The *paeA*^V321G^ mutation or *paeA* deletion was reconstructed in different *Ralstonia* GMI1000 pRalta genetic backgrounds, either *hrpG*^Q81*^ (partially intracellular infectious) (32), or *hrpG*^Q81*^-*efpR*^E66K^ (nicely intracellular infectious) (35), or *hrpG*^Q81*^-*phcQ*^R154C^ (nicely intracellular infectious) (36), or *hrcV*::Ω (extracellular infectious) (30), or *hrcS*::Ω-*vsrA*::Ω (partially intracellular infectious) (49). At 15 dpi, plants inoculated with the mutants (**A**) or the natural symbiont *C. taiwanensis* (**B**) were incubated with an excess of acetylene for four hours. Ethylene produced was measured by gas chromatography. Areas of ethylene peaks were integrated and normalized by the number of nodules, the volume of gas analyzed and the time of incubation with acetylene. Red box plots correspond to the measures made with the *paeA* mutants. At least three independent experiments with three measures per experiment were performed for each strain. * Statistically different from the parental strain (*P*<0.05, pairwise Wilcoxon test).

The nitrogen-fixing capacity of *paeA* mutants is not specific to the *hrpG*-*efpR* background as we could also measure nitrogenase activity in nodules induced by a *hrpG*-*phcQ-ΔpaeA* mutant, another genetic background conferring a good level of intracellular infection in nodules (36). In contrast, no nitrogenase activity was detected in nodules induced by *hrcV*-Δ*paeA* or *hrcS*-*vsrA-*Δ*paeA* or *hrpG-paeA*^V321G^ mutants (Fig. 3A), three genetic backgrounds known to confer only extracellular or partially intracellular infection capacity (32, 49).

Notably, the levels of nitrogenase activity detected in *hrpG*-*efpR* or *hrpG*-*phcQ* backgrounds are very low, representing less than 1% of the levels measured at 15 dpi with the natural symbiont *Cupriavidus taiwanensis* LMG19424 (Fig. 3B), and do not support plant growth (Fig. S4). Moreover, this activity is transient as it decreased at 21 dpi (Fig. S5).

### Inactivation of *paeA* increases bacterial proliferation in nodules

To determine whether the observed nitrogenase activity resulted from increased bacterial proliferation in nodules, we quantified the number of viable bacteria recovered from nodules induced by *paeA* mutants reconstructed in different genetic backgrounds. These backgrounds, *hrpG*, *hrpG-efpR*, *hrpG-phcQ*, *hrcV*, and *hrcS-vsrA*, represent the various evolutionary contexts in which *paeA* mutations arose. We found that both the *paeA*^V321G^ point mutation and the deletion of *paeA* significantly enhanced bacterial proliferation within nodules across all tested backgrounds (by a factor of 3 to 8), with the exception of the *hrcS-vsrA* background (Fig. 4A).

**Fig. 4.**
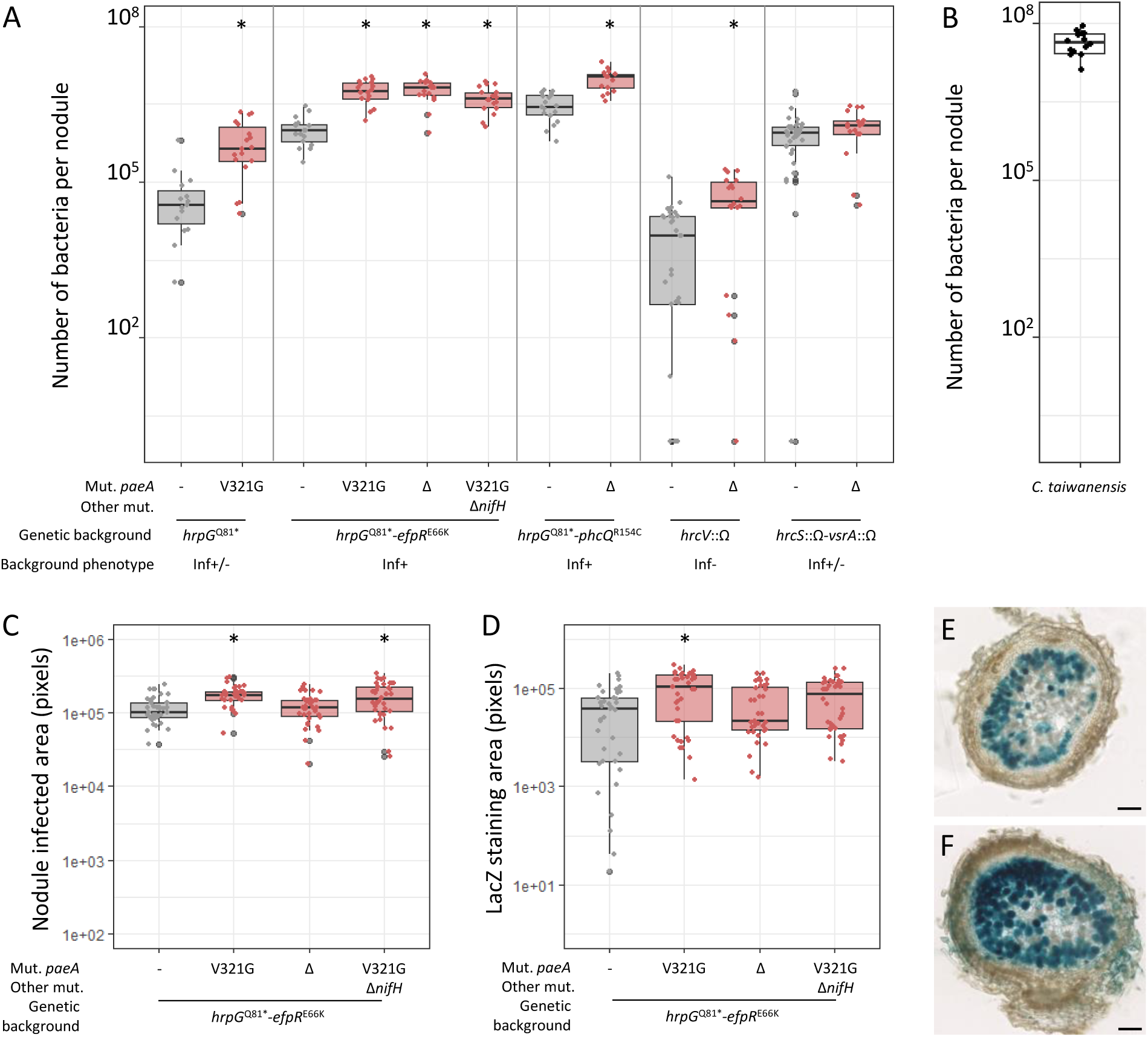
Nodule infection of *paeA* reconstructed mutants. The *paeA*^V321G^ mutation and *paeA* deletion (Δ) were reconstructed in different *Ralstonia* GMI1000 pRalta genetic backgrounds, *hrpG*^Q81*^, *hrpG*^Q81*^-*efpR*^E66K^, *hrpG*^Q81*^-*efpR*^E66K^-Δ*nifH*, *hrpG*^Q81*^-*phcQ*^R154C^, *hrcV*::Ω, or *hrcS*::Ω-*vsrA*::Ω. (**A**) Proliferation of *paeA* mutants and parental strains in 15 day-old nodules. (**B**) Proliferation of *C. taiwanensis* in 15 days-old nodules. (**C**) Total intracellularly infected areas (blue-stained + brawn-stained plant cells) and (**D**) blue-stained areas measured on 60 µm sections of 15 day-old nodules induced by *paeA* mutants and the corresponding parental strain constitutively expressing the *lacZ* gene. (**E,F**) LacZ-stained representative nodules formed by the *hrpG*^Q81*^-*efpR*^E66K^ (**E**) and *hrpG*^Q81*^-*efpR*^E66K^-*paeA*^V321G^ (**F**) mutants, both expressing constitutively the *lacZ* gene. Bars represent 100 µm length. (**A,B,C,D,E,F**) Data are from at least three independent experiments. (**A,C,D**) Red box plots correspond to measures made with the *paeA* mutants. *Statistically different from the parental strain (*P*<0.05, pairwise Wilcoxon test).

The highest numbers of bacteria recovered per nodule were observed when *paeA* mutations were combined with other highly infective mutations, such as *efpR*^E66K^ or *phcQ*^R154C^, the same genetic backgrounds in which nitrogenase activity was detected. Complementation of the *paeA*^V321G^ mutant with the wild-type allele in the *hrpG-efpR* background reduced bacterial proliferation in nodules (Fig. S3B). Furthermore, inactivation of the *nifH* gene in the *hrpG-efpR-paeA* mutant did not significantly affect the number of viable bacteria per nodule (Fig. 4A). These results suggest that nitrogenase activity is not driving the enhanced proliferation of *paeA* mutants. Rather, we propose the opposite: increased bacterial numbers in nodules may enhance the detectability of nitrogenase activity that would otherwise remain below detection thresholds.

We next investigated whether the higher number of viable bacteria recovered per nodule could be attributed to improved bacterial survival within nodule cells. To explore this, we performed live/dead staining on nodule sections. At 15 days post-inoculation (dpi), no significant differences were observed between nodules induced by the *paeA*^V321G^ mutant and those induced by its parental *hrpG-efpR* strain. By this time point, bacterial degeneration had already begun in both types of nodules (Fig. S6).

As an alternative approach, we used strains constitutively expressing the *lacZ* gene to quantify nodule cells containing the LacZ protein, whose persistence may outlast early stages of bacterial degeneration. Nodule sections were stained with X-gal, and we quantified both the area of blue-stained cells (invaded by bacteria still expressing LacZ) and brown-stained cells (invaded by bacteria that had ceased expressing LacZ). At 15 dpi, the blue-stained area was slightly larger in nodules induced by the *paeA*^V321G^ mutant (Fig. 4D–F), suggesting a modest enhancement in bacterial survival. Additionally, the total infected area per nodule section (blue + brown cells) was marginally larger in nodules induced by the mutant, indicating a slight increase in the number of cells invaded (Fig. 4C). However, these effects were minor and were not observed in the *paeA* deletion mutant.

Overall, these results suggest that *paeA* mutations exert several additive effects, including increased bacterial proliferation within nodule cells, a slightly greater number of invaded cells, and slightly improved bacterial survival. Collectively, these factors contribute to a fivefold increase, on average, in the number of viable bacteria recovered per nodule compared to nodules induced by the corresponding parental strains.

### PaeA inactivation impairs putrescine export

To determine whether the *R. pseudosolanacearum* PaeA is involved in polyamine export, like in *Salmonella* Typhimurium and *E. coli* (44), we analysed culture supernatants from parental strains and *paeA* mutants reconstructed in two genetic backgrounds, *hrpG* and *hrpG-efpR*, using mass spectrometry. The strains were grown in a synthetic medium with glutamine as the sole carbon source. In this medium, no growth difference was observed between the *paeA* mutants and the corresponding parental strains (Table S1). Consistent with previous findings, we confirmed that *R. pseudosolanacearum* GMI1000 pRalta, like GMI1000, produces very high amounts of putrescine in this medium (42, 46, 48). These levels are slightly increased in the *hrpG*-*efpR* mutant compared to the simple *hrpG* mutant. In contrast, we observed that putrescine levels were extremely low in the supernatants of *paeA* mutants in both the *hrpG* and *hrpG-efpR* genetic contexts. Complementation of the *paeA* mutants with the wild-type allele fully restored putrescine secretion (Fig. 5), confirming the major role of this protein in putrescine export. It is also noteworthy that the level of secreted putrescine in the natural symbiont *C. taiwanensis* was also very low when cultured in the same medium, despite the presence of a putrescine biosynthesis gene (RALTA_A2412) and a close homolog of *paeA* (RALTA_A0755) in the *C. taiwanensis* LMG19424 genome (Fig. S1A).

**Fig. 5.**
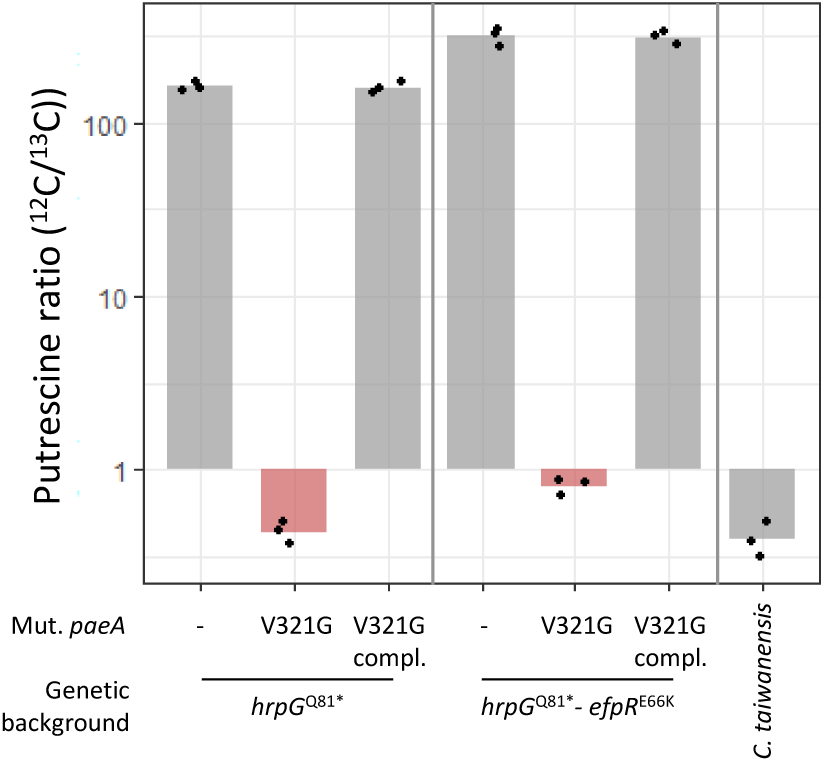
Quantification of putrescine in culture supernatants. The *R. pseudosolanacearum paeA* mutants reconstructed in GMI1000 pRalta *hrpG*^Q81*^ or GMI1000 pRalta *hrpG*^Q81*^-*efpR*^E66K^ backgrounds, the corresponding parental and complemented strains and *C. taiwanensis* were grown in minimal medium containing 10 mM glutamine until an optical density at 600 nm equal to 1. The supernatants were separated from the cells by centrifugation and then filtered at 0.2 µm in order to quantify only extracellular putrescine. Quantification of putrescine was carried out by high-resolution mass spectrometry by calculating the ratio between putrescine in the sample and the internal standard (full ^13^C-labelled putrescine). Compl. complementation of the *paeA*^V321G^ mutant with *paeA* wild-type allele introduced at the native locus. Data are from three independent experiments.

### Inactivation of *paeA* in *Ralstonia* symbionts modulates plant gene expression towards a more functional symbiotic state

In a previous study using both *R. pseudosolanacearum* and *C. taiwanensis* strains, we identified *Mimosa* genes whose expression in nodules correlated with the adaptation level of the symbionts (43). Among these genes, the putrescine biosynthesis gene encoding an arginine decarboxylase (ADC) was found to be highly expressed in *C. taiwanensis*-infected nodules, whereas it showed only weak expression in nodules infected with the *R. pseudosolanacearum* GMI1000 *pRalta hrpG efpR* mutant. Similarly, genes essential for functional nodules, such as leghemoglobins, exhibited low expression in nodules infected by *R. pseudosolanacearum* but high expression in nodules infected by *C. taiwanensis*. Additionally, we identified *Mimosa* genes whose expression was negatively correlated with bacterial adaptation to symbiosis. For instance, genes putatively involved in defense responses, such as pathogenesis-related proteins of class 10 (PR10) (51) and peroxidases (52, 53), as well as some genes associated with gibberellin biosynthesis, a phytohormone known to be finely regulated during nodule development (54), were progressively downregulated.

To further investigate the correlation between the expression of these plant genes and bacterial adaptation, we analyzed their expression by qRT-PCR in nodules induced by the *R. pseudosolanacearum hrpG-efpR-paeA* mutant. We compared these results to nodules induced by the *R. pseudosolanacearum hrpG* and *hrpG-efpR* mutants, as well as to nodules induced by *C. taiwanensis*, including either the wild-type strain or a nitrogen-fixation-deficient *nifH* mutant. Nodules were harvested at 10 dpi, an early stage chosen to ensure that *Ralstonia* bacteria had not yet significantly degenerated or had only just begun to do so. The expression of these genes in nodules induced by the *hrpG-efpR-paeA* mutant differed only slightly from their expression in nodules induced by the *hrpG-efpR* mutant. However, expression levels consistently fell between those observed in nodules induced by the *hrpG-efpR* mutant and those in nodules induced by the *C. taiwanensis* non-fixing mutant (Fig. 6). Specifically, the putrescine biosynthesis gene ADC and two globin-encoding genes exhibited increased expression, whereas PR10, a peroxidase, and a gibberellin 3-beta dioxygenase encoding gene were down-regulated in nodules induced by the *hrpG-efpR-paeA* mutant compared to nodules induced by the *hrpG-efpR* mutant. These data indicate that *R. pseudosolanacearum* symbionts carrying mutations in *paeA* triggered plant gene expression changes, shifting nodules towards a more functional symbiotic state.

**Fig. 6.**
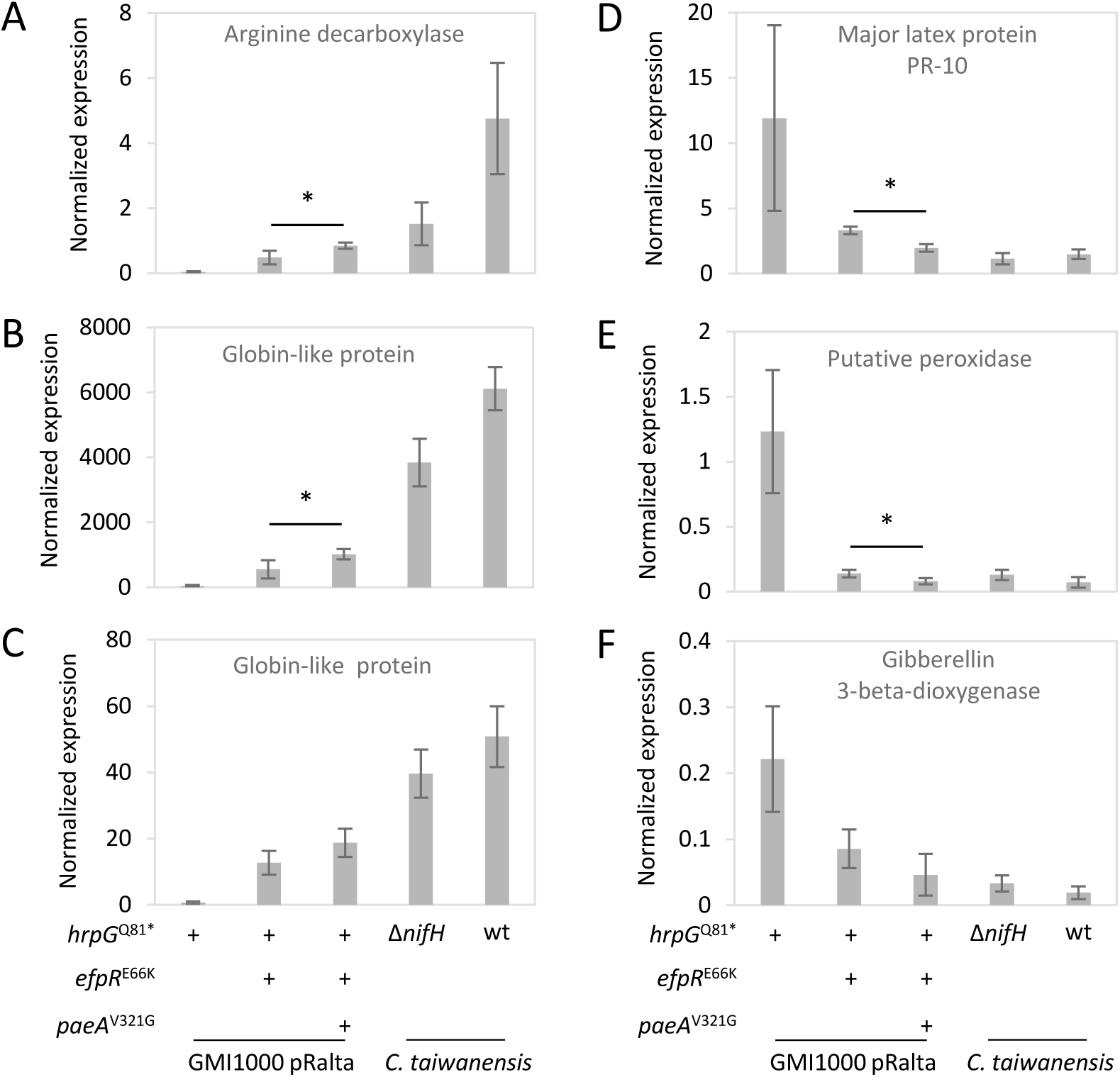
Gene expression of *Mimosa pudica* in response to progressively adapted *R. pseudosolanacaerum* symbionts and *C. taiwanensis* strains. Expression of *M. pudica* genes encoding an arginine decarboxylase (MpudA1P6v1r1_Scf09g0369161) (**A**), two globin-like proteins (MpudA1P6v1r1_Scf07g0334541 (**B**) and MpudA1P6v1r1_Scf29g0605241 (**C**), a major latex protein PR-10 (MpudA1P6v1r1_Scf15g0080361) (**D**), a putative peroxidase (MpudA1P6v1r1_Scf04g0255631) (**E**) and a gibberellin 3-beta-dioxygenase (MpudA1P6v1r1_Scf32g0613491) (**F**) was measured in 10 day-old nodules by qRT-PCR. Gene expressions were normalized by three reference genes encoding a putative ubiquitin protein (MpudA1P6v1r1_Scf06g0302791) and two putative DNA helicases (MpudA1P6v1r1_Scf39g0686881 and MpudA1P6v1r1_Scf10g0050861). wt, wild-type strain of *C. taiwanensis*. Δ*nifH*, non-fixing strain of *C. taiwanensis* deleted in the *nifH* gene. Data represent mean values and standard deviations from 4 independent experiments. * Significant difference between nodules induced by *hrpG*^Q81*^*-efpR^E66K^* and *hrpG*^Q81*^*-efpR^E66K^-paeA^V321G^* mutants (*P*<0.05, Student t-test).

## DISCUSSION

In this study, we identified in *Ralstonia pseudosolanacearum* GMI1000 a putrescine exporter, PaeA (RSc2277), whose inactivation enhances the symbiotic interaction between this bacterium and the legume *Mimosa pudica*.

Putrescine, along with spermine and spermidine, belongs to a class of small polycationic molecules known as polyamines, which are found across all three domains of life (55). Through their electrostatic interactions with various anionic macromolecules, polyamines play a crucial role in fundamental cellular processes, including protein synthesis by facilitating ribosomal subunit assembly and activity, DNA replication and chromosomal stability, as well as membrane and cell wall integrity (56–58). Beyond these essential physiological functions, polyamines have increasingly been recognized as key players in bacterial virulence. Their role in host-pathogen interactions is complex, as both hosts and microbes can synthesize polyamines and exploit them for their own benefit (59, 60). Hosts produce polyamines as part of their defense response, primarily through the generation of reactive oxygen species (ROS) from polyamine oxidation (61). Conversely, pathogens accumulate polyamines to counteract oxidative stress and enhance their survival within the host (62, 63). Additionally, pathogens utilize polyamines to regulate the expression of virulence factors. For example, putrescine enhances the production of plant cell wall-degrading enzymes and upregulates genes involved in chemotaxis and flagellar biogenesis in the plant pathogen *Dickeya fangzhongdai* (64). In other plant pathogens (*Dickeya zeae*) and human pathogens (*Proteus mirabilis*, *Yersinia pestis*), putrescine acts as an extracellular signal to modulate bacterial motility and biofilm formation, two functions important for host cell invasion (65–68). Interestingly, some pathogens also exploit host-derived polyamines for infection. For instance, *Salmonella typhimurium* and *Pseudomonas aeruginosa* uptake host-produced spermidine, which enables the expression and assembly of type III secretion systems (T3SS) essential for bacterial infection (69, 70).

In *R. pseudosolanacearum*, putrescine biosynthesis is essential, as a mutant lacking *speC* (RSc2365), which encodes an ornithine decarboxylase, is unable to grow in the absence of exogenous putrescine (46). Moreover, this bacterium excretes large amounts of putrescine in the xylem sap of host plants as well as in synthetic culture media, whereas only trace amounts of cadaverine and no detectable spermidine have been observed (46, 48). Additionally, *R. pseudosolanacearum* was shown to boost the synthesis of host putrescine during its interaction with tomato through the secretion of a type III transcription-activator-like effector (TALE) called Bgr11 that activates the expression of the tomato *ADC* gene encoding an arginine decarboxylase involved in putrescine biosynthesis. Effector-mediated activation of this gene leads to elevated agmatine and putrescine levels in tomato root and leaf tissues (71). The function of this host- and microbe-produced putrescine *in planta* remains unclear. Wu et al. (71) showed that the increased ADC activity on the plant side did not affect the growth of *R. pseudosolanacearum* in leaves.

In a previous study, we experimentally evolved *R. pseudosolanacearum* into a legume symbiont capable of forming nodules on the roots of *Mimosa pudica* and infecting these nodules intracellularly. Nodulation and intracellular infection were mainly achieved through a combination of mutations that inactivated the regulators *hrpG* and *efpR* (35) or *hrpG* and components of the *PhcA* regulatory pathway (36). Here, we found that mutations in the putrescine exporter *paeA* occurred independently six times during this evolution experiment. Repeated mutations in the same gene are usually a strong indication that these mutations confer a fitness advantage to bacteria. We demonstrated that the V321G mutation in *paeA* completely abolished putrescine export in synthetic medium and significantly enhanced bacterial proliferation in nodules, resulting in an approximately five-fold increase in viable bacteria recovered per nodule. This increase in bacterial proliferation was observed in most of the genetic backgrounds tested (GMI1000 pRalta *hrcV*, *hrpG*, *hrpG*-*efpR*, and *hrpG*-*phcQ*), indicating that the infection phenotype does not depend on the presence of specific mutations other than those conferring nodulation, either *hrcV* or *hrpG*. Furthermore, the additive effect of the *paeA* mutation on nodule infection, when combined with another highly adaptive mutation for intracellular infection (such as *efpR*^E66K^ or *phcQ*^R154C^), allowed the detection of nitrogenase activity in nodules, that would otherwise have been too low to be observed. No higher levels of nitrogenase activity were detected in nodules formed by evolved clones of this evolution experiment. Moreover, nitrogenase activity in *hrpG-efpR-paeA*-induced nodules was only transiently detected at 15 dpi and then decreased to very low level at 21 dpi. This transient phenotype likely reflects the limited intracellular persistence of evolved symbionts. Although the *paeA* mutation seems to slightly improve intracellular persistence, this trait is far from being optimized in experimentally evolved *R. pseudosolanacearum* symbionts compared to natural rhizobia. Even after 60 serial evolution cycles on *M. pudica*. *R. pseudosolanacearum* did not acquire the ability to survive for long in the cytoplasm of nodule cells, a trait found almost exclusively in rhizobia among plant-associated bacteria (10) and which may require specific functions (72).

One hypothesis to explain the improved adaptation of the *paeA* mutants to symbiosis with legumes is that the absence of polyamine secretion makes the strain intrinsically more fit. However, no growth differences were observed between the mutants and their respective parental strains in synthetic medium containing glutamine as carbon source, a medium in which *R. pseudosolanacearum* secretes large amounts of putrescine. An alternative hypothesis is that the putrescine produced by bacteria negatively interferes with the symbiotic process. Interestingly, the natural symbiont *C. taiwanensis* also does not secrete putrescine in synthetic medium containing glutamine as carbon source. Several similarities between natural and experimental evolutionary processes have already been highlighted in this experiment (73). This finding provides another striking example of parallel between the two processes, demonstrating the relevance of experimental evolution approaches for understanding natural evolution (74).

Another interesting finding is that the expression profile of *Mimosa* genes gradually changes as bacteria progressively adapt to symbiosis (43). In nodules induced by the *hrpG-efpR-paeA* mutant, the expression of certain genes approaches the levels measured in nodules induced by the natural symbiont *C. taiwanensis*. Specifically, several defense-related genes are down-regulated, reinforcing the idea that polyamines excreted by the bacterium are perceived negatively by the plant. In contrast, the expression of other plant genes essential for functional symbiosis, such as genes encoding leghemoglobins, is slightly increased in nodules induced by the *hrpG-efpR-paeA* mutant. Although the expression of these genes remains much lower than in *C. taiwanensis*-induced nodules, it indicates that bacterial evolution is progressing towards a more effective symbiosis. An intriguing observation is the progressive up-regulation of the plant ADC gene as the bacteria adapt to the symbiosis. This gene encodes a key enzyme of the putrescine biosynthetic pathway in plants. Its up-regulation is not due to the action of the type III TALE effector Bgr11 (71) in nodules since the T3SS of *R. pseudosolancearum* is inactivated in nodulating strains (32). Moreover, this up-regulation is not specifically correlated with the presence of *paeA* mutated symbionts. An increase in its expression was also observed between *hrpG*-induced and *hrpG-efpR*-induced nodules, with further up-regulation in nodules formed by the *hrpG-efpR-paeA* mutant and even more by *C. taiwanensis* symbionts. Instead, plant putrescine biosynthesis may be correlated with nodule development. Polyamine levels in legume nodules have been reported to be five to ten times higher than in other plant organs, though their composition varies considerably among legume species and appears to be an intrinsic characteristic of each species (75). The physiological role of putrescine in nodules is not entirely clear, but in *Vigna* nodules, polyamine levels, particularly putrescine, correlate linearly with both nitrogenase activity and leghemoglobin levels (76). Additionally, a study in *Lotus japonicus* suggested that polyamines primarily contribute to cell division and expansion during nodule development (77). Once again, the changes in *Mimosa ADC* gene expression in nodules induced by the *hrpG-efpR-paeA* mutant support the idea of an evolutionary trajectory towards better-developed nodules. These results also suggest that polyamines play a complex role in plant-microbe interactions and that bacterial and plant-derived putrescine contribute differently to nodule symbiosis.

In conclusion, our study highlights the role of extracellular putrescine in the adaptation of *Ralstonia pseudosolanacearum* to symbiosis with *Mimosa pudica*. The inactivation of the putrescine exporter *paeA* contributes to a shift towards a more advanced symbiotic state. While the mutation does not confer full symbiotic competence comparable to natural rhizobia, it represents a step towards bacterial adaptation by modulating host responses and improving bacterial proliferation within nodules giving the first signs of nitrogen fixation. Despite these advances, *R. pseudosolanacearum* remains limited in its ability to persist intracellularly and sustain nitrogen fixation over time. This highlights the challenges of evolving a non-rhizobial species into a mutualistic nitrogen-fixing symbiont and suggests that additional genetic changes are required. Future studies should focus on identifying the key rhizobial determinants that mediate this evolutionary transition.

## MATERIALS AND METHODS

### Bacterial strains and growth conditions

Bacterial strains used in this study are listed in Table S2. *Ralstonia pseudosolanacearum* derived strains were grown at 28°C either on rich Phi medium containing 10 g.L^-1^ bacto-peptone, 1 g.L^-1^ yeast extract and 1 g.L^-1^ casamino acids (78) or on synthetic medium (25 mM KH_2_PO_4_, 3.8 mM (NH_4_)_2_SO_4_, 0.203 mM MgSO_4_.7H_2_O, 40 µM Na_2_EDTA.2H_2_O, 15.6 µM ZnSO_4_.7H_2_O, 1.26 µM CoCl_2_.6H_2_O, 5 µM MnCl_2_.4H_2_O, 16.1 µM H_3_BO_3_, 1.6 µM Na_2_MoO_4_.2H_2_O, 10.8 µM FeSO_4_.7H_2_O, 1.2 µM CuSO_4_.5H_2_O) supplemented with 2% glycerol for natural transformation or 10 mM glutamine for putrescine quantification. The *Cupriavidus taiwanensis* strain was grown at 28°C on rich TY medium (tryptone 5 g.L^-1^, yeast extract 3 g.L^-1^). *E. coli* strains were grown in LB medium at 37°C. Antibiotics were used at the following concentrations: trimethoprim 100 μg.mL^-1^, spectinomycin 40 μg.mL^-1^, kanamycin 50 μg.mL^-1^ for *R. pseudosolanacearum* and 25 µg.mL^-1^ for *E. coli*, tetracycline 10 μg.mL^-1^, gentamicin 10 µg.mL^-1^.

### Plant material

*Mimosa pudica* seeds (LIPME production obtained from one commercial seed [B&T World Seed, Paguignan, France] of Australian origin) were sterilized as described (34). Then seedlings were transferred in glass tubes (two seedlings per tube) in N-free conditions, containing a Fahraeus slant agar and liquid Jensen ¼ medium (79, 80). Plants were grown at 28°C in a growth chamber under the following conditions: 16 h light and 8 h dark with 70% humidity.

### Experimental evolution

The evolution experiment was conducted as previously described (34, 81). Five lineages, two (B and F) derived from the CBM212 ancestor, two (G and K) derived from the CBM349 ancestor, and one (M) derived from the CBM356 ancestor, previously evolved for 35 cycles (34), were further evolved until cycle 60 using 21-day cycles of nodulation. A new lineage (X) was derived from the reconstructed mutant GMI1000 pRalta *hrpG*^Q81*^ *efpR*^E66K^ and evolved for 15 nodulation cycles of 21-days.

### Bacterial proliferation in nodules

After 15 days post inoculation, 5 to 10 nodules per plant from 6 different plants per strain analyzed were collected independently, sterilized in 2.4% hypochlorite solution for 15 min, rinsed three times with sterile H_2_O and ground in 1 mL sterile H_2_O. Serial dilutions were plated on Phi medium containing trimethoprim and grown for 48 h at 28°C.

### Acetylene reduction assays

At 15 or 21 days post inoculation, three sets of 6 plants were collected for each inoculated strain. The number of nodules was counted and the 6 plants were transferred together to 60 mL test tubes sealed with a Suba-seal^Ⓡ^septa. A syringe was used to remove 1 ml of air and replace it with 1 ml of acetylene. Plants were incubated for 4 hours at 28°C in the light. The ethylene produced was measured by analyzing 0.4 mL of gas using an Agilent 7820A gas chromatograph system with GS-Alumina column (ref. 115-3552). The area of the ethylene peak was integrated and normalized by the number of nodules, the volume of gas analyzed and the number of hours of incubation with acetylene.

### Cytological analyses

Nodules were harvested at 15 days post inoculation and fixed in Z’ buffer (potassium phosphate buffer 0.1 M, KCl 10 mM, MgSO_4_ 1 mM, pH7.4) with 2.5% glutaraldehyde and 0.1% triton for 1 h under vacuum. Nodules were embedded in 4% agarose and sectioned at 60 µm using a Leica VT1000S vibratome. Sections were incubated in a staining solution (potassium ferricyanide and ferrocyanide 5 mM each, 0.008% XGal in Z’ buffer) for 1 h at 37°C. The largest longitudinal section was observed using an axioplan microscope (Zeiss) or a NanoZoomer scanner (Hamamatsu). The blue areas revealed by LacZ staining and brown areas corresponding to nodule invaded cells were measured in pixels using the Fiji software with the HSB color model. Statistical significance was analyzed using the pairwise Wilcoxon test. Live dead staining of nodule sections were performed using Live/dead Baclight Bacterial Viability Kit (Invitrogen L7012). Sections were incubated 15 min at room temperature in the dark. The largest longitudinal section was observed by an inverted fluorescence microscope Nikon Eclipse Ti.

### Quantification of extracellular putrescine

Bacterial strains were grown in a synthetic medium containing 10 mM glutamine as sole carbon source at 28°C until an optical density at 600 nm (OD_600_) of 1 (∼5×10^8^ CFU.mL^-1^). Two milliliters of bacterial cultures were centrifuged for 5 min at maximum speed. Then, the supernatants were filtered using a syringe equipped with a 0.2 µm filter. The filtered supernatants were diluted at 1/10 with ultra pure water and then diluted at ½ with IDMS (Isotopic Dilution Mass Spectrometry). Finally, samples were transferred in HPLC vials and analyzed by liquid chromatography-mass spectrometry using an UHPLC vanquish (Thermo Fisher), with HS F5 DISCOVERY 150 x 2.1 mm i.d., particle size 5 µm (Supelco) column with guard column SUPELGUARD KIT HS F5 5µm 20 X2.1 mm (Supelco) at 30°C. The mobile phases were water (A) and acetonitrile (B) both containing 0.1% of formic acid. The LC gradient program was: 0 min to 15 min, 0% B; 30 min, 35% B; 35 min to 40 min, 40% B; 40 min to 45 min, 0% B. The injection volume was 5 μL. The HPLC were coupled to a detection by high resolute ion mass spectrometry with an Orbitrap exploris 120 orbitrap (Thermo Fischer). Full scan HRMS analyses were performed in FTMS mode at a resolution of 60 000 (at 200 m/z), with the following source parameters: Spray voltage: 3400 V, ion transfer tube temperature: 320°C, vaporization temperature: 75°C, sheath gas 25 arb, Auxiliary gas 5 arb. Data processing were performed by skyline (version 24.1).

Other materials and methods concerning genome resequencing and detection of mutations in evolved clones, constructions of mutants and plasmids, phylogeny of evolved clones of the B lineage and *M. pudica* gene expression analyses by quantitative reverse transcription-PCRs are provided in Text S1 in the supplemental material.

## ACKNOWLEDGEMENTS

This study was supported by the French National Research Agency (ANR-22-CE20-0014 and ANR-21-CE02-0019-01), the “Laboratoires d’Excellence (LABEX)” TULIP (ANR-10-LABX-41), and the “École Universitaire de Recherche (EUR)” TULIP-GS (ANR-18-EURE-0019). This work was performed in collaboration with the GeT core facility, Toulouse, France (GeT, https://doi.org/10.15454/1.5572370921303193E12). GeT core facility was supported by France Génomique National infrastructure, funded as part of “Investissement d’avenir” program managed by Agence Nationale pour la Recherche (contract ANR-10-INBS-09). We thank Cécile Pouzet, Yves Martinez and Aurélie Leru for their help with the cytological analyses at the TRI FR3450 cellular imaging facility, Castanet-Tolosan, France.

## SUPPLEMENTAL MATERIALS

Fig S1. Protein sequence and 3D structure of *R. pseudosolanacearum* PaeA

Fig. S2. Phylogeny of evolved clones of the B lineage.

Fig. S3. Complementation of the *paeA*^V321G^ mutant in the *hrpG*^Q81*^*-efpR*^E66K^ genetic background

Fig. S4. Effect of *Ralstonia paeA* mutants on plant growth.

Fig. S5. Acetylene reduction assays at 15 and 21 days post-inoculation.

Fig S6. Intracellular persistence of bacteroids evaluated by LIVE/DEAD staining.

Table S1. Growth rate of the *paeA* mutants and their parental strains in synthetic medium containing 10 mM glutamine as carbon source

Table S2. Strains and plasmids used in this study.

Table S3. Mutations present in the evolved clones represented in Fig. 2.

Table S4. Primers used in this study.

Text S1. Supplemental materials and methods

## Notes

### Competing Interest Statement

The authors have declared no competing interest.

## REFERENCES

1. Limpens E, Franken C, Smit P, Willemse J, Bisseling T, Geurts R. 2003. LysM domain receptor kinases regulating rhizobial Nod factor-induced infection. Science 302:630–3. 10.1126/science.1090074.

2. Madsen EB, Madsen LH, Radutoiu S, Olbryt M, Rakwalska M, Szczyglowski K, Sato S, Kaneko T, Tabata S, Sandal N, Stougaard J. 2003. A receptor kinase gene of the LysM type is involved in legume perception of rhizobial signals. Nature 425:637–40. 10.1038/nature02045.

3. Radutoiu S, Madsen LH, Madsen EB, Felle HH, Umehara Y, Grønlund M, Sato S, Nakamura Y, Tabata S, Sandal N, Stougaard J. 2003. Plant recognition of symbiotic bacteria requires two LysM receptor-like kinases. Nature 425:585–92. 10.1038/nature02039.

4. Arrighi JF, Barre A, Ben Amor B, Bersoult A, Soriano LC, Mirabella R, de Carvalho-Niebel F, Journet EP, Ghérardi M, Huguet T, Geurts R, Dénarié J, Rougé P, Gough C. 2006. The *Medicago truncatula* lysin motif-receptor-like kinase gene family includes NFP and new nodule-expressed genes. Plant Physiol 142:265–79. 10.1104/pp.106.084657.

5. Smit P, Limpens E, Geurts R, Fedorova E, Dolgikh E, Gough C, Bisseling T. 2007. Medicago LYK3, an entry receptor in rhizobial nodulation factor signaling. Plant Physiol 145:183–91. 10.1104/pp.107.100495.

6. Gage DJ. 2002. Analysis of infection thread development using Gfp- and DsRed-expressing *Sinorhizobium meliloti*. J Bacteriol 184:7042–7046. 10.1128/JB.184.24.7042-7046.2002.

7. Sprent JI. 2007. Evolving ideas of legume evolution and diversity: a taxonomic perspective on the occurrence of nodulation. New Phytologist 174:11–25. 10.1111/j.1469-8137.2007.02015.x.

8. Montiel J, Reid D, Grønbæk TH, Benfeldt CM, James EK, Ott T, Ditengou FA, Nadzieja M, Kelly S, Stougaard J. 2021. Distinct signaling routes mediate intercellular and intracellular rhizobial infection in *Lotus japonicus*. Plant Physiol 185:1131–1147. 10.1093/plphys/kiaa049.

9. Coba de la Peña T, Fedorova E, Pueyo JJ, Lucas MM. 2017. The symbiosome: legume and rhizobia co-evolution toward a nitrogen-fixing organelle? Front Plant Sci 8:2229. 10.3389/fpls.2017.02229.

10. Parniske M. 2018. Uptake of bacteria into living plant cells, the unifying and distinct feature of the nitrogen-fixing root nodule symbiosis. Curr Opin Plant Biol 44:164–174. 10.1016/j.pbi.2018.05.016.

11. Masson-Boivin C, Sachs JL. 2018. Symbiotic nitrogen fixation by rhizobia-the roots of a success story. Curr Opin Plant Biol 44:7–15. 10.1016/j.pbi.2017.12.001.

12. Downie JA. 2010. The roles of extracellular proteins, polysaccharides and signals in the interactions of rhizobia with legume roots. FEMS Microbiol Rev 34:150–170. 10.1111/j.1574-6976.2009.00205.x.

13. López-Baena FJ, Ruiz-Sainz JE, Rodríguez-Carvajal MA, Vinardell JM. 2016. Bacterial molecular signals in the *Sinorhizobium fredii*-soybean symbiosis. Int J Mol Sci 17. 10.3390/ijms17050755.

14. Oono R, Schmitt I, Sprent JI, Denison RF. 2010. Multiple evolutionary origins of legume traits leading to extreme rhizobial differentiation. New Phytologist 187:508–520. 10.1111/j.1469-8137.2010.03261.x.

15. Czernic P, Gully D, Cartieaux F, Moulin L, Guefrachi I, Patrel D, Pierre O, Fardoux J, Chaintreuil C, Nguyen P, Gressent F, Da Silva C, Poulain J, Wincker P, Rofidal V, Hem S, Barrière Q, Arrighi JF, Mergaert P, Giraud E. 2015. Convergent evolution of endosymbiont differentiation in Dalbergioid and Inverted Repeat-Lacking Clade legumes mediated by nodule-specific cysteine-rich peptides. Plant Physiol 169:1254–65. 10.1104/pp.15.00584.

16. Mergaert P. 2018. Role of antimicrobial peptides in controlling symbiotic bacterial populations. Nat Prod Rep 35:336–356. 10.1039/c7np00056a.

17. Farkas A, Maróti G, Durgő H, Györgypál Z, Lima RM, Medzihradszky KF, Kereszt A, Mergaert P, Kondorosi É. 2014. *Medicago truncatula* symbiotic peptide NCR247 contributes to bacteroid differentiation through multiple mechanisms. Proc Natl Acad Sci U S A 111:5183–8. 10.1073/pnas.1404169111.

18. Penterman J, Abo RP, De Nisco NJ, Arnold MF, Longhi R, Zanda M, Walker GC. 2014. Host plant peptides elicit a transcriptional response to control the *Sinorhizobium meliloti* cell cycle during symbiosis. Proc Natl Acad Sci U S A 111:3561–6. 10.1073/pnas.1400450111.

19. Sankari S, Babu VMP, Bian K, Alhhazmi A, Andorfer MC, Avalos DM, Smith TA, Yoon K, Drennan CL, Yaffe MB, Lourido S, Walker GC. 2022. A haem-sequestering plant peptide promotes iron uptake in symbiotic bacteria. Nat Microbiol 7:1453–1465. 10.1038/s41564-022-01192-y.

20. Horváth B, Domonkos Á, Kereszt A, Szűcs A, Ábrahám E, Ayaydin F, Bóka K, Chen Y, Chen R, Murray JD, Udvardi MK, Kondorosi É, Kaló P. 2015. Loss of the nodule-specific cysteine rich peptide, NCR169, abolishes symbiotic nitrogen fixation in the *Medicago truncatula dnf7* mutant. Proc Natl Acad Sci U S A 112:15232–7. 10.1073/pnas.1500777112.

21. Horváth B, Güngör B, Tóth M, Domonkos Á, Ayaydin F, Saifi F, Chen Y, Biró JB, Bourge M, Szabó Z, Tóth Z, Chen R, Kaló P. 2023. The *Medicago truncatula* nodule-specific cysteine-rich peptides, NCR343 and NCR-new35 are required for the maintenance of rhizobia in nitrogen-fixing nodules. New Phytol 239:1974–1988. 10.1111/nph.19097.

22. Marlow VL, Haag AF, Kobayashi H, Fletcher V, Scocchi M, Walker GC, Ferguson GP. 2009. Essential role for the BacA protein in the uptake of a truncated eukaryotic peptide in *Sinorhizobium meliloti*. J Bacteriol 191:1519–1527. 10.1128/JB.01661-08.

23. Guefrachi I, Pierre O, Timchenko T, Alunni B, Barrière Q, Czernic P, Villaécija-Aguilar JA, Verly C, Bourge M, Fardoux J, Mars M, Kondorosi E, Giraud E, Mergaert P. 2015. *Bradyrhizobium* BclA is a peptide transporter required for bacterial differentiation in symbiosis with *Aeschynomene* legumes. Mol Plant Microbe Interact 28:1155–66. 10.1094/MPMI-04-15-0094-R.

24. Glazebrook J, Ichige A, Walker GC. 1993. A *Rhizobium meliloti* homolog of the *Escherichia coli* peptide-antibiotic transport protein SbmA is essential for bacteroid development. Genes Dev 7:1485–97. 10.1101/gad.7.8.1485.

25. Crespo-Rivas JC, Guefrachi I, Mok KC, Villaécija-Aguilar JA, Acosta-Jurado S, Pierre O, Ruiz-Sainz JE, Taga ME, Mergaert P, Vinardell JM. 2016. *Sinorhizobium fredii* HH103 bacteroids are not terminally differentiated and show altered O-antigen in nodules of the Inverted Repeat-Lacking Clade legume *Glycyrrhiza uralensis*. Environ Microbiol 18:2392–404. 10.1111/1462-2920.13101.

26. Barrière Q, Guefrachi I, Gully D, Lamouche F, Pierre O, Fardoux J, Chaintreuil C, Alunni B, Timchenko T, Giraud E, Mergaert P. 2017. Integrated roles of BclA and DD-carboxypeptidase 1 in *Bradyrhizobium* differentiation within NCR-producing and NCR-lacking root nodules. Sci Rep 7:9063. 10.1038/s41598-017-08830-0.

27. Chen WF, Wang ET, Ji ZJ, Zhang JJ. 2021. Recent development and new insight of diversification and symbiosis specificity of legume rhizobia: mechanism and application. J Appl Microbiol 131:553–563. 10.1111/jam.14960.

28. Hirsch AM, Wilson KJ, Jones JD, Bang M, Walker VV, Ausubel FM. 1984. *Rhizobium meliloti* nodulation genes allow *Agrobacterium tumefaciens* and *Escherichia coli* to form pseudonodules on alfalfa. J Bacteriol 158:1133–43. 10.1128/jb.158.3.1133-1143.1984.

29. Abe M, Kawamura R, Higashi S, Mori S, Shibata M, Uchiumi T. 1998. Transfer of the symbiotic plasmid from *Rhizobium leguminosarum* biovar *trifolii* to *Agrobacterium tumefaciens*. J Gen Appl Microbiol 44:65–74. 10.2323/jgam.44.65.

30. Nakatsukasa H, Uchiumi T, Kucho K, Suzuki A, Higashi S, Abe M. 2008. Transposon mediation allows a symbiotic plasmid of *Rhizobium leguminosarum* bv. *trifolii* to become a symbiosis island in *Agrobacterium and Rhizobium*. J Gen Appl Microbiol 54:107–18. 10.2323/jgam.54.107.

31. Nandasena KG, O’Hara GW, Tiwari RP, Sezmis E, Howieson JG. 2007. *In situ* lateral transfer of symbiosis islands results in rapid evolution of diverse competitive strains of mesorhizobia suboptimal in symbiotic nitrogen fixation on the pasture legume *Biserrula pelecinus* L. Environ Microbiol 9:2496–2511. 10.1111/j.1462-2920.2007.01368.x.

32. Marchetti M, Capela D, Glew M, Cruveiller S, Chane-Woon-Ming B, Gris C, Timmers T, Poinsot V, Gilbert LB, Heeb P, Medigue C, Batut J, Masson-Boivin C. 2010. Experimental evolution of a plant pathogen into a legume symbiont. PLoS Biol 8 (1):e1000280. 10.1371/journal.pbio.1000280

33. Valls M, Genin S, Boucher C. 2006. Integrated regulation of the type III secretion system and other virulence determinants in *Ralstonia solanacearum*. PLoS Pathog 2:798–807. 10.1371/journal.ppat.0020082.

34. Doin de Moura GG, Mouffok S, Gaudu N, Cazalé AC, Milhes M, Bulach T, Valière S, Roche D, Ferdy JB, Masson-Boivin C, Capela D, Remigi P. 2023. A selective bottleneck during host entry drives the evolution of new legume symbionts. Mol Biol Evol 40. 10.1093/molbev/msad116.

35. Capela D, Marchetti M, Clérissi C, Perrier A, Guetta D, Gris C, Valls M, Jauneau A, Cruveiller S, Rocha EPC, Masson-Boivin C. 2017. Recruitment of a lineage-specific virulence regulatory pathway promotes intracellular infection by a plant pathogen experimentally evolved into a legume symbiont. Mol Biol Evol 34:2503–2521. 10.1093/molbev/msx165.

36. Tang M, Bouchez O, Cruveiller S, Masson-Boivin C, Capela D. 2020. Modulation of quorum sensing as an adaptation to nodule cell infection during experimental evolution of legume symbionts. mBio 11(1):e03129–19. 10.1128/mbio.03129-19

37. Perrier A, Peyraud R, Rengel D, Barlet X, Lucasson E, Gouzy J, Peeters N, Genin S, Guidot A. 2016. Enhanced *in planta* fitness through adaptive mutations in EfpR, a dual regulator of virulence and metabolic functions in the plant pathogen *Ralstonia solanacearum*. PLoS Pathog 12:e1006044. 10.1371/journal.ppat.1006044.

38. Khokhani D, Lowe-Power TM, Tran TM, Allen C. 2017. A single regulator mediates strategic switching between attachment/spread and growth/virulence in the plant pathogen. mBio 8(5):e00895–17. 10.1128/mbio.00895-17

39. Mori Y, Ishikawa S, Ohnishi H, Shimatani M, Morikawa Y, Hayashi K, Ohnishi K, Kiba A, Kai K, Hikichi Y. 2017. Involvement of ralfuranones in the quorum sensing signalling pathway and virulence of *Ralstonia solanacearum* strain OE1-1. Mol Plant Pathol. 10.1111/mpp.12537.

40. Perrier A, Barlet X, Peyraud R, Rengel D, Guidot A, Genin S. 2018. Comparative transcriptomic studies identify specific expression patterns of virulence factors under the control of the master regulator PhcA in the *Ralstonia solanacearum* species complex. Microb Pathog 116:273–278. 10.1016/j.micpath.2018.01.028.

41. Takemura C, Senuma W, Hayashi K, Minami A, Terazawa Y, Kaneoka C, Sakata M, Chen M, Zhang Y, Nobori T, Sato M, Kiba A, Ohnishi K, Tsuda K, Kai K, Hikichi Y. 2021. PhcQ mainly contributes to the regulation of quorum sensing-dependent genes, in which PhcR is partially involved, in *Ralstonia pseudosolanacearum* strain OE1-1. Mol Plant Pathol 22:1538–1552. 10.1111/mpp.13124.

42. Peyraud R, Cottret L, Marmiesse L, Gouzy J, Genin S. 2016. A resource allocation trade-off between virulence and proliferation drives metabolic versatility in the plant pathogen *Ralstonia solanacearum*. PLoS Pathog 12:e1005939. 10.1371/journal.ppat.1005939.

43. Libourel C, Keller J, Brichet L, Cazalé AC, Carrère S, Vernié T, Couzigou JM, Callot C, Dufau I, Cauet S, Marande W, Bulach T, Suin A, Masson-Boivin C, Remigi P, Delaux PM, Capela D. 2023. Comparative phylotranscriptomics reveals ancestral and derived root nodule symbiosis programmes. Nat Plants 9:1067–1080. 10.1038/s41477-023-01441-w.

44. Iwadate Y, Ramezanifard R, Golubeva YA, Fenlon LA, Slauch JM. 2021. PaeA (YtfL) protects from cadaverine and putrescine stress in *Salmonella Typhimurium* and *E. coli*. Mol Microbiol 115:1379–1394. 10.1111/mmi.14686.

45. Iwadate Y, Golubeva YA, Slauch JM. 2023. Cation homeostasis: coordinate regulation of polyamine and magnesium levels in *Salmonella*. mBio 14:e0269822. 10.1128/mbio.02698-22.

46. Lowe-Power TM, Hendrich CG, von Roepenack-Lahaye E, Li B, Wu D, Mitra R, Dalsing BL, Ricca P, Naidoo J, Cook D, Jancewicz A, Masson P, Thomma B, Lahaye T, Michael AJ, Allen C. 2018. Metabolomics of tomato xylem sap during bacterial wilt reveals *Ralstonia solanacearum* produces abundant putrescine, a metabolite that accelerates wilt disease. Environ Microbiol 20:1330–1349. 10.1111/1462-2920.14020.

47. Gerlin L, Escourrou A, Cassan C, Maviane Macia F, Peeters N, Genin S, Baroukh C. 2021. Unravelling physiological signatures of tomato bacterial wilt and xylem metabolites exploited by *Ralstonia solanacearum*. Environ Microbiol 23:5962–5978. 10.1111/1462-2920.15535.

48. Baroukh C, Zemouri M, Genin S. 2022. Trophic preferences of the pathogen *Ralstonia solanacearum* and consequences on its growth in xylem sap. Microbiologyopen 11:e1240. 10.1002/mbo3.1240.

49. Guan SH, Gris C, Cruveiller S, Pouzet C, Tasse L, Leru A, Maillard A, Medigue C, Batut J, Masson-Boivin C, Capela D. 2013. Experimental evolution of nodule intracellular infection in legume symbionts. ISME J 7:1367–1377. 10.1038/ismej.2013.24.

50. Remigi P, Capela D, Clerissi C, Tasse L, Torchet R, Bouchez O, Batut J, Cruveiller S, Rocha EP, Masson-Boivin C. 2014. Transient hypermutagenesis accelerates the evolution of legume endosymbionts following horizontal gene transfer. PLoS Biol 12:e1001942. 10.1371/journal.pbio.1001942.

51. Lopes NDS, Santos AS, de Novais DPS, Pirovani CP, Micheli F. 2023. Pathogenesis-related protein 10 in resistance to biotic stress: progress in elucidating functions, regulation and modes of action. Front Plant Sci 14:1193873. 10.3389/fpls.2023.1193873.

52. Almagro L, Gómez Ros LV, Belchi-Navarro S, Bru R, Ros Barceló A, Pedreño MA. 2009. Class III peroxidases in plant defence reactions. J Exp Bot 60:377–90. 10.1093/jxb/ern277.

53. Freitas CDT, Costa JH, Germano TA, de O Rocha R, Ramos MV, Bezerra LP. 2024. Class III plant peroxidases: from classification to physiological functions. Int J Biol Macromol 263:130306. 10.1016/j.ijbiomac.2024.130306.

54. Lin J, Frank M, Reid D. 2020. No home without hormones: how plant hormones control legume nodule organogenesis. Plant Commun 1:100104. 10.1016/j.xplc.2020.100104.

55. Michael AJ. 2016. Polyamines in eukaryotes, bacteria, and archaea. J Biol Chem 291:14896–903. 10.1074/jbc.R116.734780.

56. Tabor CW, Tabor H. 1985. Polyamines in microorganisms. Microbiol Rev 49:81–99. 10.1128/mr.49.1.81-99.1985.

57. Igarashi K, Kashiwagi K. 2018. Effects of polyamines on protein synthesis and growth of *Escherichia coli*. J Biol Chem 293:18702–18709. 10.1074/jbc.TM118.003465.

58. Duprey A, Groisman EA. 2020. DNA supercoiling differences in bacteria result from disparate DNA gyrase activation by polyamines. PLoS Genet 16:e1009085. 10.1371/journal.pgen.1009085.

59. Gerlin L, Baroukh C, Genin S. 2021. Polyamines: double agents in disease and plant immunity. Trends Plant Sci 26:1061–1071. 10.1016/j.tplants.2021.05.007.

60. Yi Q, Park MJ, Vo KTX, Jeon JS. 2024. Polyamines in plant-pathogen interactions: roles in defense mechanisms and pathogenicity with applications in fungicide development. Int J Mol Sci 25. 10.3390/ijms252010927.

61. Blázquez MA. 2024. Polyamines: their role in plant development and stress. Annu Rev Plant Biol 75:95–117. 10.1146/annurev-arplant-070623-110056.

62. Campilongo R, Di Martino ML, Marcocci L, Pietrangeli P, Leuzzi A, Grossi M, Casalino M, Nicoletti M, Micheli G, Colonna B, Prosseda G. 2014. Molecular and functional profiling of the polyamine content in enteroinvasive *E. coli*: looking into the gap between commensal *E. coli* and harmful *Shigella*. PLoS One 9:e106589. 10.1371/journal.pone.0106589.

63. Nair AV, Singh A, Rajmani RS, Chakravortty D. 2024. *Salmonella Typhimurium* employs spermidine to exert protection against ROS-mediated cytotoxicity and rewires host polyamine metabolism to ameliorate its survival in macrophages. Redox Biol 72:103151. 10.1016/j.redox.2024.103151.

64. Xie C, Gu W, Chen Z, Liang Z, Huang S, Zhang LH, Chen S. 2023. Polyamine signaling communications play a key role in regulating the pathogenicity of *Dickeya fangzhongdai*. Microbiol Spectr 11:e0196523. 10.1128/spectrum.01965-23.

65. Shi Z, Wang Q, Li Y, Liang Z, Xu L, Zhou J, Cui Z, Zhang LH. 2019. Putrescine is an intraspecies and interkingdom cell-cell communication signal modulating the virulence of *Dickeya zeae*. Front Microbiol 10:1950. 10.3389/fmicb.2019.01950.

66. Sturgill G, Rather PN. 2004. Evidence that putrescine acts as an extracellular signal required for swarming in Proteus mirabilis. Mol Microbiol 51:437–46. 10.1046/j.1365-2958.2003.03835.x.

67. Patel CN, Wortham BW, Lines JL, Fetherston JD, Perry RD, Oliveira MA. 2006. Polyamines are essential for the formation of plague biofilm. J Bacteriol 188:2355–63. 10.1128/JB.188.7.2355-2363.2006.

68. Kurihara S, Sakai Y, Suzuki H, Muth A, Phanstiel O, Rather PN. 2013. Putrescine importer PlaP contributes to swarming motility and urothelial cell invasion in *Proteus mirabilis*. J Biol Chem 288:15668–76. 10.1074/jbc.M113.454090.

69. Zhou L, Wang J, Zhang LH. 2007. Modulation of bacterial Type III secretion system by a spermidine transporter dependent signaling pathway. PLoS One 2:e1291. 10.1371/journal.pone.0001291.

70. Miki T, Uemura T, Kinoshita M, Ami Y, Ito M, Okada N, Furuchi T, Kurihara S, Haneda T, Minamino T, Kim YG. 2024. *Salmonella Typhimurium* exploits host polyamines for assembly of the type 3 secretion machinery. PLoS Biol 22:e3002731. 10.1371/journal.pbio.3002731.

71. Wu D, von Roepenack-Lahaye E, Buntru M, de Lange O, Schandry N, Pérez-Quintero AL, Weinberg Z, Lowe-Power TM, Szurek B, Michael AJ, Allen C, Schillberg S, Lahaye T. 2019. A plant pathogen type III effector protein subverts translational regulation to boost host polyamine levels. Cell Host Microbe 26:638–649.e5. 10.1016/j.chom.2019.09.014.

72. Fagorzi C, Ilie A, Decorosi F, Cangioli L, Viti C, Mengoni A, diCenzo GC. 2020. Symbiotic and nonsymbiotic members of the genus *Ensifer* (syn. *Sinorhizobium*) are separated into two clades based on comparative genomics and high-throughput phenotyping. Genome Biol Evol 12:2521–2534. 10.1093/gbe/evaa221.

73. Clerissi C, Touchon M, Capela D, Tang M, Cruveiller S, Parker MA, Moulin L, Masson-Boivin C, Rocha EPC. 2018. Parallels between experimental and natural evolution of legume symbionts. Nat Commun 9:2264. 10.1038/s41467-018-04778-5.

74. Stroud JT, Ratcliff WC. 2025. Long-term studies provide unique insights into evolution. Nature 639:589–601. 10.1038/s41586-025-08597-9.

75. Fujihara S, Abe H, Minakawa Y, Akao S, Yoneyama T. 1994. Polyamines in nodules from various plant-microbe symbiotic associations, vol 35, p 1127–1134. Plant and Cell Physiology. 10.1093/oxfordjournals.pcp.a078705.

76. Lahiri K, Chattopadhyay S, Ghosh B. 2004. Correlation of endogenous free polyamine levels with root nodule senescence in different genotypes in *Vigna mungo* L. J Plant Physiol 161:563–71. 10.1078/0176-1617-01057.

77. Flemetakis E, Efrose RC, Desbrosses G, Dimou M, Delis C, Aivalakis G, Udvardi MK, Katinakis P. 2004. Induction and spatial organization of polyamine biosynthesis during nodule development in *Lotus japonicus*. Mol Plant Microbe Interact 17:1283–93. 10.1094/MPMI.2004.17.12.1283.

78. Boucher CA, Barberis PA, Trigalet AP, Demery DA. 1985. Transposon mutagenesis of *Pseudomonas solanacearum* – isolation of Tn5-induced avirulent mutants. J Gen Microbiol 131:2449–2457. 10.1099/00221287-131-9-2449.

79. Fahraeus G. 1957. The infection of clover root hairs by nodule bacteria studied by a simple glass slide technique. J Gen Microbiol 16:374–81. 10.1099/00221287-16-2-374.

80. Jensen. 1942. Nitrogen fixation in leguminous plants. I. General characters of root nodule bacteria isolated from species of *Medicago* and *Trifolium* in Australia, vol 66, p 68–108, Proc. Int. Soc. N.S.W.

81. Marchetti M, Clerissi C, Yousfi Y, Gris C, Bouchez O, Rocha E, Cruveiller S, Jauneau A, Capela D, Masson-Boivin C. 2017. Experimental evolution of rhizobia may lead to either extra- or intracellular symbiotic adaptation depending on the selection regime. Mol Ecol 26:1818–1831. 10.1111/mec.13895.

