## Supplemental Figures and Tables for "Disruption of putrescine export in experimentally evolved *Ralstonia pseudosolanacearum* enhances symbiosis with *Mimosa pudica*"

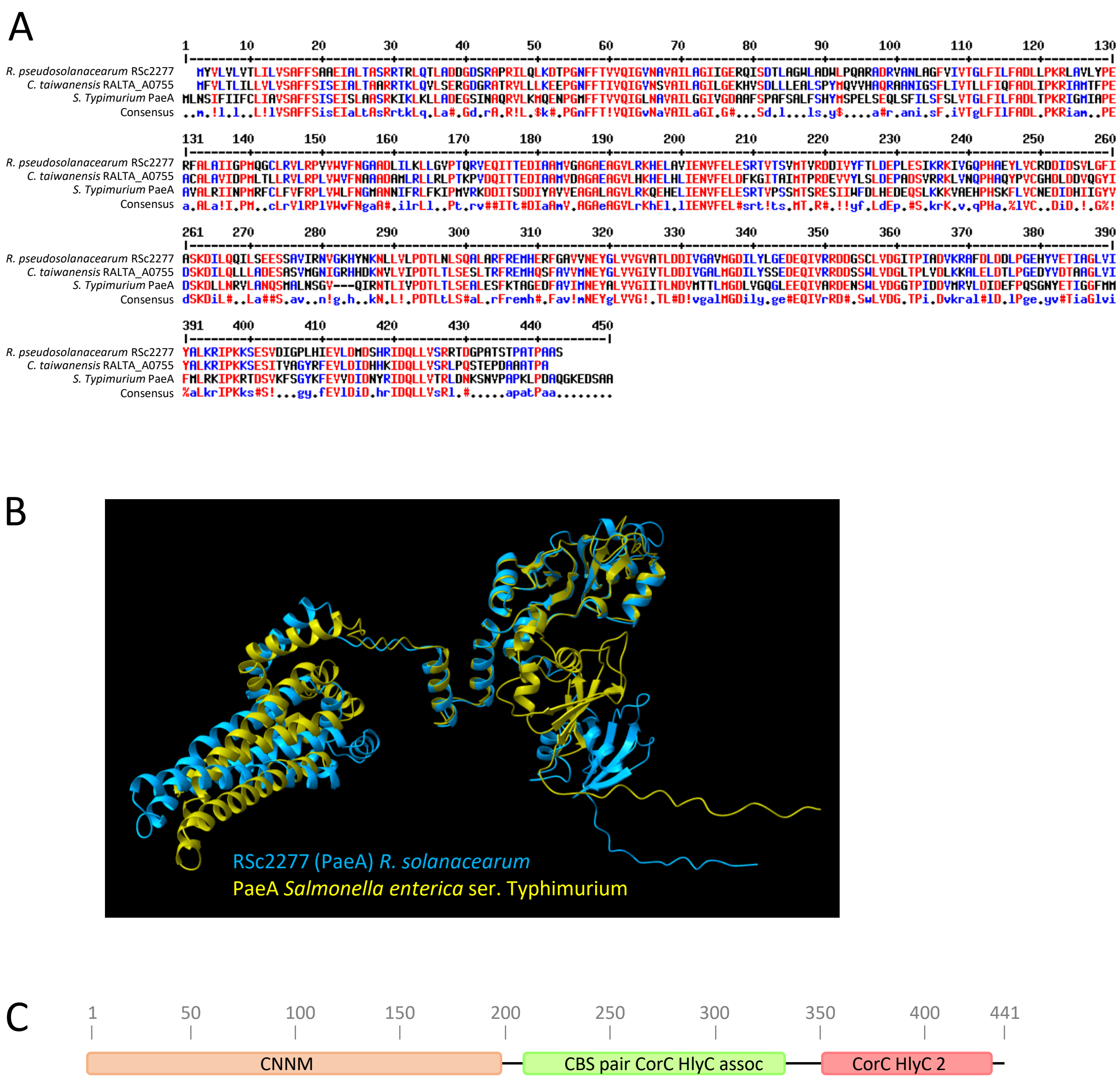


**Fig. S1. A.** Comparison of the PaeA protein sequences of *Ralstonia pseudosolanacearum* (RSc2277), *Cupriavidus taiwanensis* (RALTA_A0755) and *Salmonella enterica* ser. Typhimurium generated with Multalin (1). **B.** Three-dimensional structures of the PaeA proteins from *R. pseudosolanacearum* (in blue) and *S. enterica* ser. Typhimurium (in gold) predicted by AlphaFold (2) . Overlay generated by ChimeraX (3). The C-terminal region between amino acids 426 to 441 is a disordered region as predicted by the flDPnn software (4). **C**. InterPro scan domains of the *R. pseudosolanceartum* PaeA protein. CNNM, CBS-pair domain divalent metal cation transport mediator. CBS pair CorC HlyC assoc, two tandem repeats of the cystathionine beta-synthase (CBS pair) domains the majority of which are associated with the CorC_HlyC domain. CorC HlyC 2, domain that might be involved in modulating transport of ion substrates.

**References**

1. Corpet F. 1988. Multiple sequence alignment with hierarchical clustering. Nucleic Acids Res 16:10881-10890. https://doi.org/10.1093/nar/16.22.10881.
2. Jumper J, Evans R, Pritzel A, Green T, Figurnov M, Ronneberger O, Tunyasuvunakool K, Bates R, Žídek A, Potapenko A, Bridgland A, Meyer C, Kohl SAA, Ballard AJ, Cowie A, Romera-Paredes B, Nikolov S, Jain R, Adler J, Back T, Petersen S, Reiman D, Clancy E, Zielinski M, Steinegger M, Pacholska M, Berghammer T, Bodenstein S, Silver D, Vinyals O, Senior AW, Kavukcuoglu K, Kohli P, Hassabis D. 2021. Highly accurate protein structure prediction with AlphaFold. Nature 596:583-589. https://doi.org/10.1038/s41586-021-03819-2.
3. Meng EC, Goddard TD, Pettersen EF, Couch GS, Pearson ZJ, Morris JH, Ferrin TE. 2023. UCSF ChimeraX: Tools for structure building and analysis. Protein Sci 32:e4792. https://doi.org/10.1002/pro.4792.
4. Hu G, Katuwawala A, Wang K, Wu Z, Ghadermarzi S, Gao J, Kurgan L. 2021. flDPnn: Accurate intrinsic disorder prediction with putative propensities of disorder functions. Nat Commun 12:4438. https://doi.org/10.1038/s41467-021-24773-7.

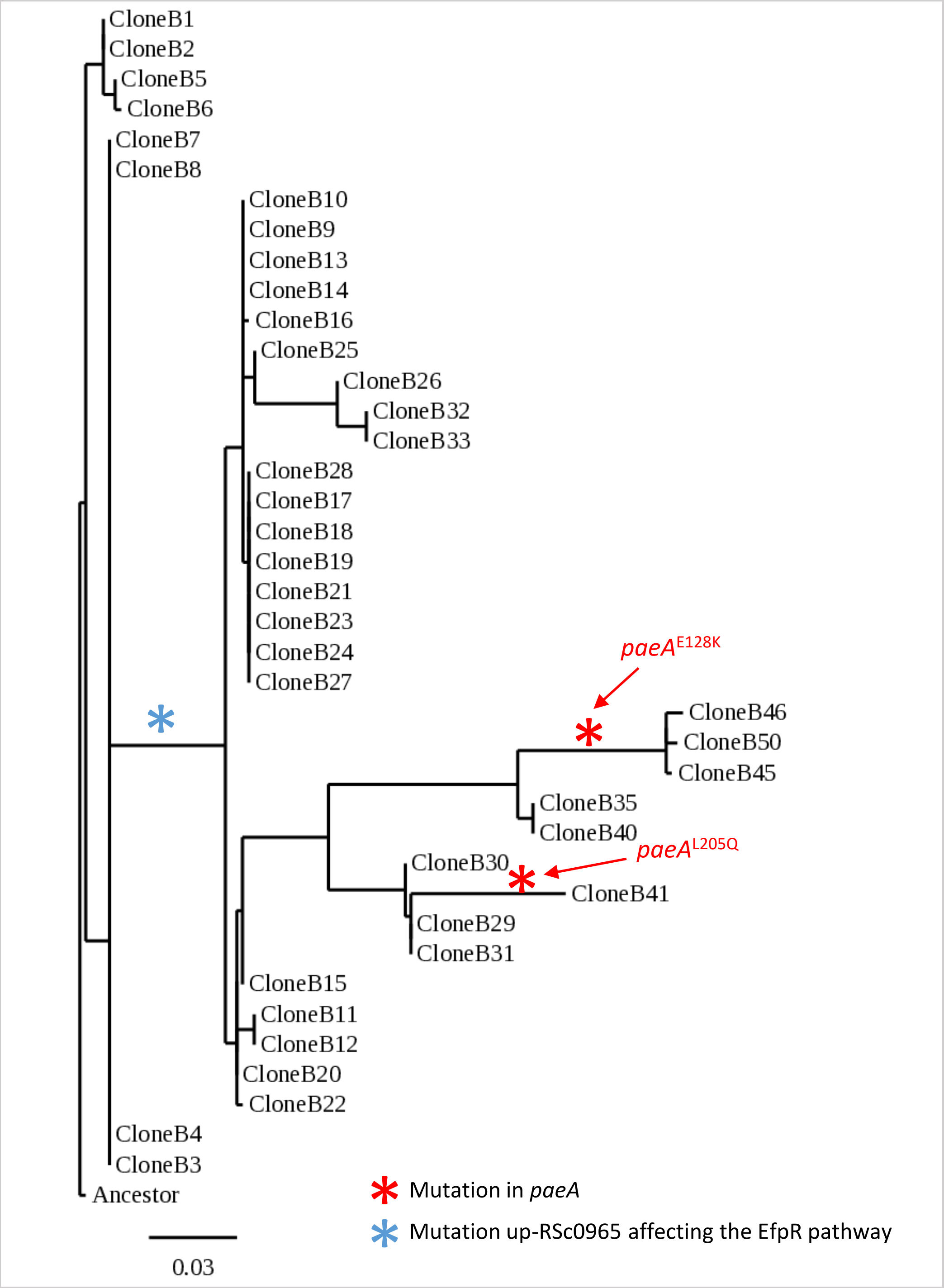


**Fig. S2. Phylogenetic tree of evolved clones from the B lineage.**
The tree was based on artificial sequences constructed by concatenating all the mutated allele positions occurring in the experiment. Maximum likelihood (ML) heuristic search under the LG model with C-rate variation among sites, as implemented in PhyMLv3.0 software, was used to construct the tree. The black bar represents the scale of genetic variation. Clone numbers indicate the cycle of the evolution experiment in which they were isolated. The blue star indicates the occurrence of the mutation up-RSc0965, which represses the EfpR regulator. The occurrence of mutations in the *paeA* gene is shown in red.

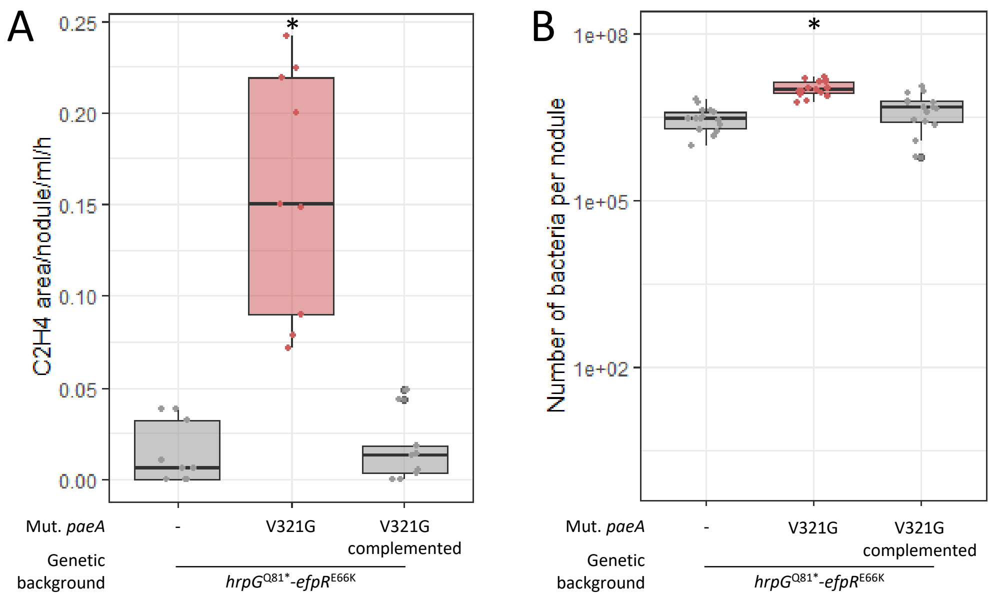


**Fig. S4. Complementation of the *paeA*$^{V321G}$ mutant in the *hrpG*$^{Q81*}$*-efpR*$^{E66K}$ genetic background**
Acetylene reduction assays (**A**) and number of viable bacteria recovered per nodule (**B**) measured on plants inoculated with the *hrpG*$^{Q81*}$-*efpR*$^{E66K}$-*paeA*$^{V321G}$ mutant (red box plot) and its parental and complemented derived strains, 15 days post inoculation. * Statistically different from the parental strain (*P*<0.05, pairwise Wilcoxon test).

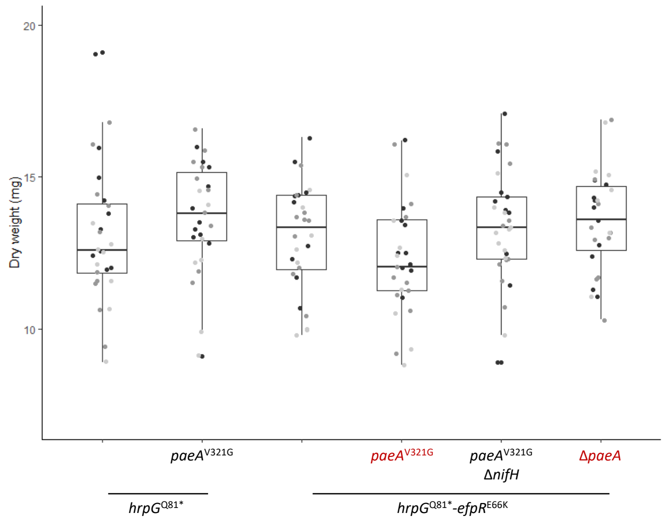


**Fig. S4. Effect of *Ralstonia paeA* mutants on plant growth.**
The aerial part of plants inoculated with *paeA* mutants in the *hrpG*$^{Q81*}$ or *hrpG*$^{Q81*}$-*efpR*$^{E66K}$ background was harvested at 28 dpi and dried at 65°C for two days. Three independent experiments were performed with 10 measurements per experiment and inoculated strain, displayed with three shades of grey. The dry weights of plants inoculated with strains showing detectable levels of nitrogenase activity (strains indicated in red) were not statistically different from the dry weights of plants inoculated with non-fixing strains (strains indicated in black) (*P*>0.05, ANOVA test).

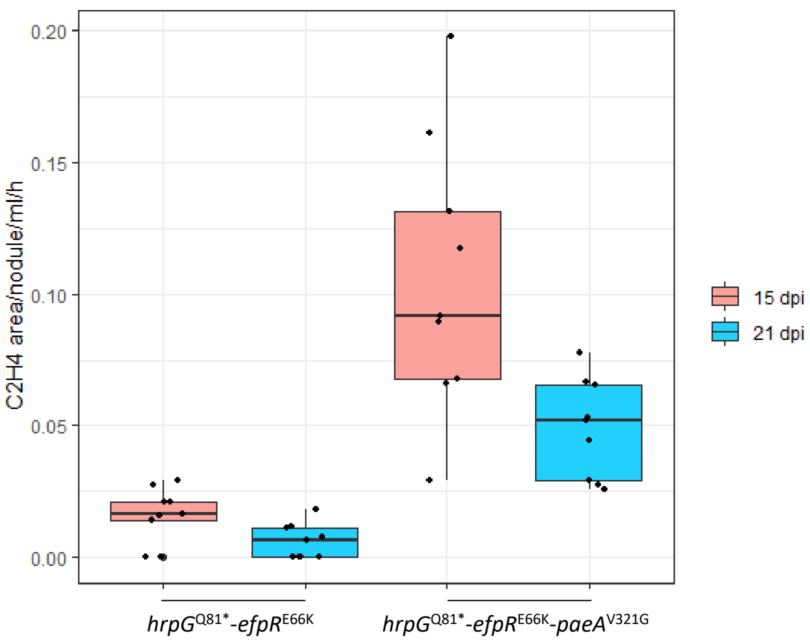


**Fig. S5. Acetylene reduction assays at 15 and 21 days post-inoculation**.
Plants were incubated with an excess of acetylene for four hours. Ethylene produced was measured by gas chromatography. Areas of ethylene peaks were integrated and normalized by the number of nodules, the volume of gas analyzed and the time of incubation with acetylene. Three independent experiments with three measures per experiment were performed for each strain.

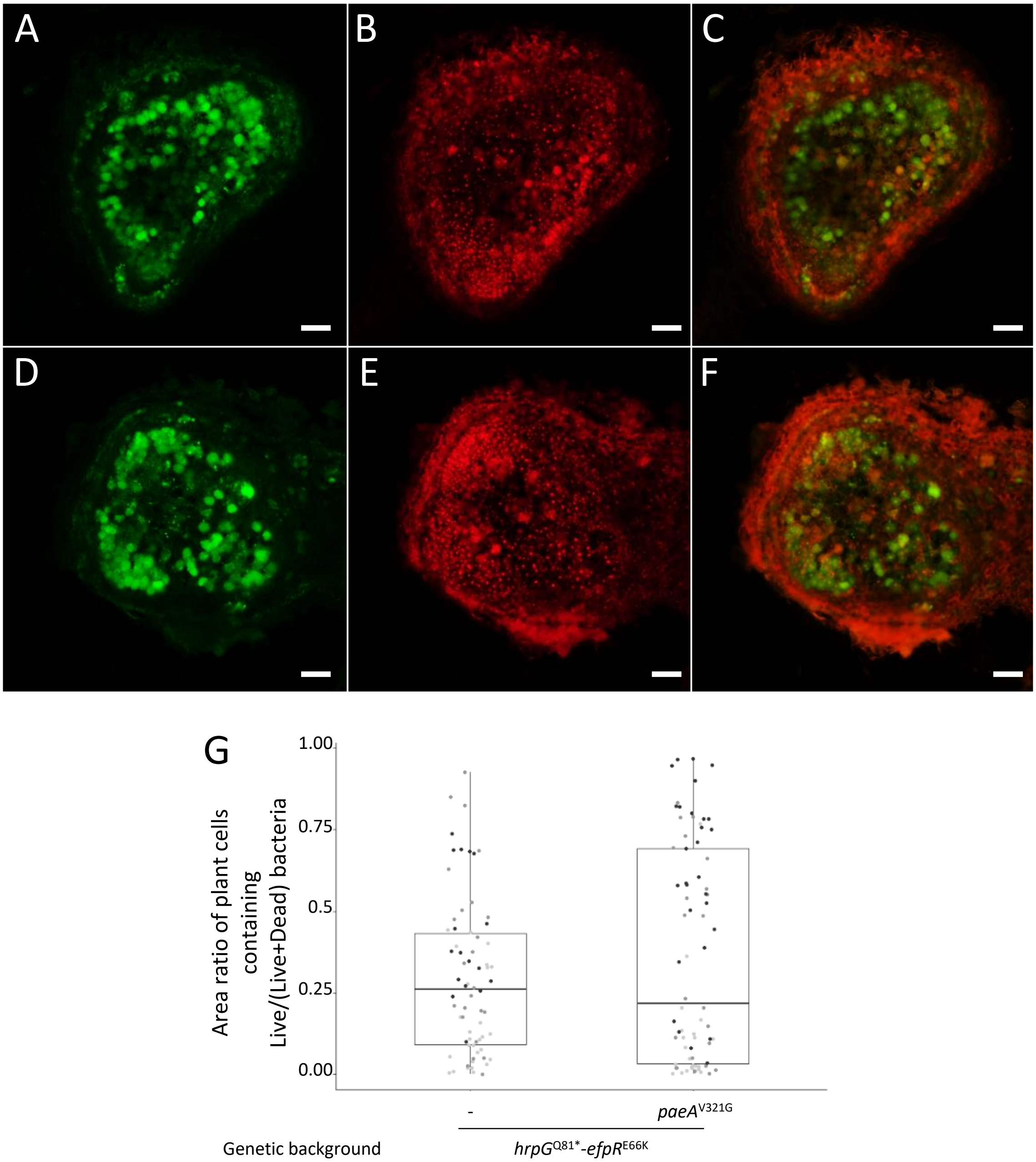


**Fig. S6**. **Intracellular persistence of bacteroids evaluated by LIVE/DEAD staining**

Sections of 15 day-old nodules formed by *R. pseudosolanacearum* GMI1000 pRalta *hrpG*$^{Q81*}$ *efpR*$^{E66K}$ (**A, B, C**) and its isogenic mutant *paeA*$^{V321G}$ (**D, E, F**) were stained with the LIVE/DEAD™ BacLight™ Bacterial Viability Kit. (**A,D**) SYTO9 staining of live cells. (**B,E**) Propidium iodide staining of dead cells. (**C,F**) Overlay of SYTO9 and propidium iodide staining. White bars represent 100 µm length. (**G**) The area ratio of plant cells containing live bacteria to (live+dead) bacteria per nodule section was evaluated. At least 16 nodules per experiment and per strain were analysed in three independent experiments, shown as grey shades. Differences were not statistically significant (*P*=0.45, Wilcoxon test).

**Table S1. Growth rates of the *paeA* mutants and their parental strains in synthetic medium containing 10 mM glutamine as carbon source**

| Strain | µmax ($h^{-1}$) | *P* value (*t*-test)* |
|---|---|---|
| GMI1000 pRalta *hrpG*$^{Q81*}$ | 0.266 ± 0.06 | 0.28459001 |
| GMI1000 pRalta *hrpG*$^{Q81*}$ *paeA*$^{V321G}$ | 0.226 ± 0.03 | |
| GMI1000 pRalta *hrpG*$^{Q81*}$ *efpR*$^{E66K}$ | 0.236 ± 0.02 | 0.10538523 |
| GMI1000 pRalta *hrpG*$^{Q81*}$ *efpR*$^{E66K}$ *paeA*$^{V321G}$ | 0.265 ± 0.02 | |

*The *t*-test was performed between the growth rates of the *paeA* mutants and the corresponding parental strains.

**Table S2.** Strains and plasmids used in this study.

| Organism<br>Strain ID | Common name | Relevant characteristics | Reference/ source |
|---|---|---|---|
| *E. coli* | | | |
| DH5α | DH5α | *F recA lacZ*DM15 | Bethesda Research Laboratory |
| *R. solanacearum* | | | |
| GMI1000 | | wild-type strain (phylotype IA) isolated from tomato in French Guyana | (1) |
| RCM3804 | GMI1000 Δ*paeA* | Introduction of an unmarked deletion of *paeA* in GMI1000 | This study |
| RCM3438 | GMI1000 *paeA*$^{V321G}$ IG*glmS*:P*psbA*-GFP | Introduction of the missense mutation *paeA*$^{V321G}$ and a P*psbA*-GFP fusion downstream *glmS* in GMI1000, KanR | This study |
| *C. taiwanensis* | | | |
| LMG19424 | | Wild-type strain isolated from *Mimosa pudica* in Taiwan | (2) |
| CBM832 | | LMG19424 derivative resistant to streptomycin, StrR | (3) |
| CBM2865 | CBM832 Δ*nifH* | Introduction of an unmarked deletion of *nifH* in CBM832, StrR | (4) |
| Chimeric *Ralstonia* | | | |
| CBM124 | GMI1000 pRalta | GMI1000 pRalta::Tri, TriR | (5) |
| CBM212 | Anc$^{B}$ | CBM124-derived nodulating ancestor of the B lineage, mutated in *hrpG*$^{Q81*}$, TriR, GenR | (5) |
| CBM349 | Anc$^{BK}$ | CBM124-derived nodulating ancestor of the G and K lineages, mutated in *hrpG*$^{Q209*}$, TriR, GenR | (5) |
| CBM356 | Anc$^{M}$ | CBM124-derived nodulating ancestor of the M lineage, mutated in *hrcV*$^{Q589*}$ (stop mutation), TriR, GenR | (5) |
| CBM1195 | B9 | Evolved clone from lineage B, cycle 9, TriR, GenR | (6) |
| RCM2305 | B30 | Evolved clone from lineage B, cycle 30, TriR, GenR | (7) |
| RCM2685 | B40 | Evolved clone from lineage B, cycle 40, TriR, GenR | This study |
| RCM2700 | B41 | Evolved clone from lineage B, cycle 41, TriR, GenR | This study |
| RCM3145 | B45 | Evolved clone from lineage B, cycle 45, TriR, GenR | This study |
| CBM465 | G2 | Evolved clone from lineage G, cycle 2, TriR, GenR | This study |
| CBM557 | G4 | Evolved clone from lineage G, cycle 4, TriR, GenR | (7) |
| CBM594 | G5 | Evolved clone from lineage G, cycle 5, TriR, GenR | (7) |
| CBM1125 | K4 | Evolved clone from lineage K, cycle 4, TriR, GenR | (7) |
| CBM1808 | K13 | Evolved clone from lineage K, cycle 13, TriR, GenR | (8), (7) |
| CBM597 | M5 | Evolved clone from lineage M, cycle 5, TriR, GenR | (6) |
| CBM643 | M6 | Evolved clone from lineage M, cycle 6, TriR, GenR | (7) |
| RCM3373 | X6 | Evolved clone from lineage X, cycle 6, TriR, KanR | This study |
| CBM1627 | GMI1000 pRalta *hrpG*$^{Q81*}$ | Introduction of the stop mutation Q81* in *hrpG* in CBM124, TriR | (9) |
| RCM3467 | GMI1000 pRalta *hrpG*$^{Q81*}$ *paeA*$^{V321G}$ IG*glmS*:P*psbA*-GFP | Introduction of the missense mutation V321G in *paeA* and a P*psbA*-GFP fusion downstream *glmS* in CBM1627, TriR, KanR | This study |
| RCM3867 | RCM3467 *paeA* wild-type IG*glmS*:P*psbA*-GFP | Complementation of the RCM3467 strain with the wild-type allele of *paeA* introduced at the same locus | This study |
| RCM1865 | GMI1000 pRalta *hrpG*$^{Q81*}$ *efpR*$^{E66K}$ IG*glmS*:KanR, Anc$^{X}$ | Introduction of the missense mutation E66K in *efpR* and a KanR cassette downstream *glmS* in CBM1627, TriR, KanR. Ancestor of the X lineage | (10) |

| | | | |
|---|---|---|---|
| RCM2940 | GMI1000 pRalta *hrpG*$^{Q81*}$ *efpR*$^{E66K}$ IG*glmS*:P*psbA-lacZ* | Exchange of the kanamycin resistance cassette by a P*psbA*-lacZ fusion downstream *glmS* in RCM1865, TriR, GenR | (11) |
| RCM3440 | GMI1000 pRalta *hrpG*$^{Q81*}$ *efpR*$^{E66K}$ *paeA*$^{V321G}$ IG*glmS*:SpeR | Introduction of the missense mutation V321G in *paeA* and a spectinomycin resistance cassette downstream *glmS* in RCM1865, TriR, KanR | This study |
| RCM3475 | GMI1000 pRalta *hrpG*$^{Q81*}$ *efpR*$^{E66K}$ *paeA*$^{V321G}$ IG*glmS*: P*psbA*-mCherry | Exchange of the spectinomycine resistance cassette by a P*psbA*-mCherry fusion downstream *glmS* in RCM3440, TriR, KanR | This study |
| RCM3642 | GMI1000 pRalta *hrpG*$^{Q81*}$ *efpR*$^{E66K}$ *paeA*$^{V321G}$ IG*glmS*:P*psbA-lacZ* | Exchange of the P*psbA*-mCherry fusion by a P*psbA-lacZ* fusion downstream *glmS* in RCM3475, TriR, KanR | This study |
| RCM3871 | RCM3440 *paeA* wild-type | Complementation of the RCM3440 strain with the wild-type allele of *paeA* introduced at the same locus | This study |
| RCM3687 | GMI1000 pRalta *hrpG*$^{Q81*}$ *efpR*$^{E66K}$ *paeA*$^{V321G}$ Δ*nifH* IG*glmS*:P*psbA*-mCherry | Introduction of an unmarked deletion of the *nifH* gene in RCM3475, TriR, KanR | This study |
| RCM3773 | GMI1000 pRalta *hrpG*$^{Q81*}$ *efpR*$^{E66K}$ Δ*paeA* IG*glmS*:KanR | Introduction of an unmarked deletion of *paeA* in RCM1865, TriR, KanR | This study |
| RCM2495 | GMI1000 pRalta *hrpG*$^{Q81*}$ *phcQ*$^{R154C}$ IG*glmS*:P*psbA*-mCherry | Introduction of the missense R154C mutation in *phcQ* and a P*psbA*-mCherry fusion downstream *glmS* in CBM1627, TriR, KanR | (8) |
| RCM3820 | GMI1000 pRalta *hrpG*$^{Q81*}$ *phcQ*$^{R154C}$ Δ*paeA* IG*glmS*:P*psbA*-mCherry | Introduction of an unmarked deletion of *paeA* in RCM2495, TriR, KanR | This study |
| CBM125 | GMI1000pRalta *hrcV::Ω* | Inactivation of *hrcV* by a spectinomycin resistance cassette in CBM124, TriR, SpeR | (5) |
| RCM3854 | GMI1000pRalta *hrcV::Ω* Δ*paeA* | Introduction of an unmarked deletion of *paeA* in CBM125, TriR, SpeR | This study |
| CBM1619 | GMI1000pRalta *hrcS::Ω vsrA::Ω* | Inactivation of *hrcS* by a kanamycin resistance cassette and *vsrA* by a spectinomycin resistance cassette in CBM124, TriR, KanR, SpeR | (12) |
| RCM3857 | GMI1000pRalta *hrcS::Ω vsrA::Ω* Δ*paeA* | Introduction of an unmarked deletion of *paeA* in CBM1619, TriR, KanR, SpeR | This study |

| **Plasmid names** | **Relevant characteristics** | **Reference** |
|---|---|---|
| pRalta | Symbiotic plasmid of LMG19424 (0.5 Mb) | (13) |
| pEX18Tc | Suicide plasmid carrying the *sacB* gene, TetR | (14) |
| pCBM142 | pGEM-T plasmid carrying the SpeR cassette inserted in the intergenic region downstream *glmS*, SpeR, AmpR | (15) |
| pRCK-P*psbA*-GFP | Plasmid for *R. solanacearum* chromosomal integration of the constitutive *psbA* promoter fused to GFPuv into the intergenic region downstream *glmS*, KanR | (10) |
| pRCK-P*psbA*-mCherry | Plasmid for *R. solanacearum* chromosomal integration of the constitutive *psbA* promoter fused to mCherry into the intergenic region downstream *glmS*, KanR | (10) |
| pRCG-P*psbA*-lacZ | Plasmid for *R. solanacearum* chromosomal integration of the constitutive *psbA* promoter fused to *lacZ* into the intergenic region downstream *glmS*, GenR | (16) |

TriR, trimethoprim resistant. SpeR, spectinomycin resistant. GenR, gentamycin resistant. KanR, kanamycin resistant. TetR, tetracyclin resistant. AmpR, ampicillin resistance.

### Supplemental references


1. Boucher CA, Barberis PA, Trigalet AP, Demery DA. 1985. Transposon mutagenesis of *Pseudomonas solanacearum* – isolation of Tn5-induced avirulent mutants. *J Gen Microbiol* 131:2449-2457.

2. Chen WM, Laevens S, Lee TM, Coenye T, De Vos P, Mergeay M, Vandamme P. 2001. *Ralstonia taiwanensis* sp nov., isolated from root nodules of *Mimosa* species and sputum of a cystic fibrosis patient. *Int J Syst Evol Microbiol* 51:1729-1735.

3. Daubech B, Poinsot V, Klonowska A, Capela D, Chaintreuil C, Moulin L, Marchetti M, Masson-Boivin C. 2019. A new nodulation gene involved in the biosynthesis of Nod Factors with an open-chain oxidized terminal residue and in the symbiosis with. *Mol Plant Microbe Interact* 32:1635-1648.

4. Daubech B, Remigi P, Doin de Moura G, Marchetti M, Pouzet C, Auriac MC, Gokhale CS, Masson-Boivin C, Capela D. 2017. Spatio-temporal control of mutualism in legumes helps spread symbiotic nitrogen fixation. Elife 6:e28683. doi: 10.7554/eLife.28683.

5. Marchetti M, Capela D, Glew M, Cruveiller S, Chane-Woon-Ming B, Gris C, Timmers T, Poinsot V, Gilbert LB, Heeb P, Medigue C, Batut J, Masson-Boivin C. 2010. Experimental evolution of a plant pathogen into a legume symbiont. *PLoS Biol* 8.

6. Marchetti M, Jauneau A, Capela D, Remigi P, Gris C, Batut J, Masson-Boivin C. 2014. Shaping bacterial symbiosis with legumes by experimental evolution. *Mol Plant Microbe Interact* 27:956-64.

7. Doin de Moura GG, Mouffok S, Gaudu N, Cazalé AC, Milhes M, Bulach T, Valière S, Roche D, Ferdy JB, Masson-Boivin C, Capela D, Remigi P. 2023. A selective bottleneck during host entry drives the evolution of new legume symbionts. *Mol Biol Evol* 40.

8. Tang M, Bouchez O, Cruveiller S, Masson-Boivin C, Capela D. 2020. Modulation of quorum sensing as an adaptation to nodule cell infection during experimental evolution of legume symbionts. *mBio* 11.

9. Guan SH, Gris C, Cruveiller S, Pouzet C, Tasse L, Leru A, Maillard A, Médigue C, Batut J, Masson-Boivin C, Capela D. 2013. Experimental evolution of nodule intracellular infection in legume symbionts. *ISME J* 7:1367-77.

10. Capela D, Marchetti M, Clérissi C, Perrier A, Guetta D, Gris C, Valls M, Jauneau A, Cruveiller S, Rocha EPC, Masson-Boivin C. 2017. Recruitment of a lineage-specific virulence regulatory pathway promotes intracellular infection by a plant pathogen experimentally evolved into a legume symbiont. *Mol Biol Evol* 34:2503-2521.

11. Libourel C, Keller J, Brichet L, Cazalé AC, Carrère S, Vernié T, Couzigou JM, Callot C, Dufau I, Cauet S, Marande W, Bulach T, Suin A, Masson-Boivin C, Remigi P, Delaux PM, Capela D. 2023. Comparative phylotranscriptomics reveals ancestral and derived root nodule symbiosis programmes. *Nat Plants* 9:1067-1080.

12. Guan SH, Gris C, Cruveiller S, Pouzet C, Tasse L, Leru A, Maillard A, Medigue C, Batut J, Masson-Boivin C, Capela D. 2013. Experimental evolution of nodule intracellular infection in legume symbionts. *ISME J* 7:1367-1377.

13. Amadou C, Pascal G, Mangenot S, Glew M, Bontemps C, Capela D, Carrere S, Cruveiller S, Dossat C, Lajus A, Marchetti M, Poinsot V, Rouy Z, Servin B, Saad M, Schenowitz C, Barbe V, Batut J, Medigue C, Masson-Boivin C. 2008. Genome sequence of the beta-rhizobium *Cupriavidus taiwanensis* and comparative genomics of rhizobia. *Genome Res* 18:1472-1483.

14. Hoang TT, Karkhoff-Schweizer RR, Kutchma AJ, Schweizer HP. 1998. A broad-host-range Flp-FRT recombination system for site-specific excision of chromosomally-located DNA sequences: application for isolation of unmarked *Pseudomonas aeruginosa* mutants. *Gene* 212:77-86.

15. Remigi P, Capela D, Clerissi C, Tasse L, Torchet R, Bouchez O, Batut J, Cruveiller S, Rocha EP, Masson-Boivin C. 2014. Transient hypermutagenesis accelerates the evolution of legume endosymbionts following horizontal gene transfer. *PLoS Biol* 12:e1001942.

16. Monteiro F, Solé M, van Dijk I, Valls M. 2012. A chromosomal insertion toolbox for promoter probing, mutant complementation, and pathogenicity studies in *Ralstonia solanacearum*. *Mol Plant Microbe Interact* 25:557-68.

**Table S4. Primers used in this study**

| Forward primer name | Forward primer sequence 5'-3' | Reverse primer name | Reverse primer sequence 5'-3' | Amplified region | Product length |
|---|---|---|---|---|---|
| oCBM4077 | CTCCTGCATCAGCACTTC | oCBM4078 | GGATGGACGCGTACATTC | Amplification of 6 kb fragment around the mutation V321G position 2474088 in paeA | 6085 bp |
| oCBM4079 | GCGCGACGACATCGTCTACTTC | oCBM4080 | TCGACGCTCTCGCTCTTCTTGG | Amplification of a fragment around the mutation V321G position 2474088 in paeA for sequencing the region | 558 bp |
| oCBM4339 | CAACGAATACGGGCTGGTGCT | oCBM4080 | TCGACGCTCTCGCTCTTCTTGG | Screening oligos for detecting the wild-type allele of paeA (sequence position 2474088) | 263 bp |
| oCBM4340 | CAACGAATACGGGCTGGTGCG | oCBM4080 | TCGACGCTCTCGCTCTTCTTGG | Screening oligos for detecting the mutant allele of paeA (sequence position 2474088) | 263 bp |
| oCBM5016 | CCCAAGCTTCGATAGCCCTTGTCCCACAG | oCBM5017 | GCTCTAGAGGATCAGCGTGACCAGAAC | Amplification of the region upstream of paeA for cloning into the pEX18Tc plasmid to delete the gene (HindIII and XbaI overhang restriction sites) | 942 bp |
| oCBM5018 | GCTCTAGATCCTGAGCGGCGAACATG | oCBM5019 | CGGGATCCGCCGCATGTGACTTTCTGAC | Amplification of the region downstream of paeA for cloning into the pEX18Tc plasmid to delete the gene (BamHI and XbaI overhang restriction sites) | 748 bp |
| oCBM5020 | CTGCGGCAGGATCTCGAAG | oCBM5021 | CGTTGAATCGTTGCGGTGC | Amplification around the deleted paeA region to verify paeA deletion mutants | 3192 bp if the region is wild-type / 1904 bp if the region is deleted |
| oCBM4957 | CCCAAGCTTAGCTGCTATCCGGATGTTCG | oCBM4958 | GCTCTAGATCCTCAGTCTGCGGTGTTTC | Amplification of the region upstream of nifH for cloning into the pEX18Tc plasmid to delete the gene (HindIII and XbaI overhang restriction sites) | 797 bp |
| oCBM4959 | GCTCTAGATGGACTACGCGATGACCAC | oCBM4960 | CGGGATCCGGGTGTTCTCTAGTTCCGCC | Amplification of the region downstream of nifH for cloning into the pEX18Tc plasmid to delete the gene (BamHI and XbaI overhang restriction sites) | 817 bp |
| oCBM5012 | TTCCAGTGACTGTCGAAGCT | oCBM5013 | TAGTTCCGCCAATGTACCGT | Amplification around the deleted nifH region to verify nifH deletion mutants | 2751 bp if the region is wild-type / 1737 bp if the region is deleted |
| oCBM5014 | TCATCGTTGGGTGTGATCCA | oCBM5015 | GTTCTGCATGCTGGACTACG | Amplification of an internal fragment of nifH to verify nifH deletion mutants | 561 bp if the gene is wild-type / 0 bp if the gene is deleted |
| oCBM5687 | TGAAGACCGTGCCTCATTCC | oCBM5688 | TAGAGGGAGGCGGACATTGA | Amplification of a fragment of the Mimosa pudica MpudA1P6v1r1_Scf09g0369161 endocing an arginine decarboxylase for qPCR | 127 bp |
| oCBM5022 | TGGTGAAAGAGGCACTGTTG | oCBM5023 | TATGCTTTGGCCCATGCATTG | Amplification of a fragment of the Mimosa pudica MpudA1P6v1r1_Scf07g0334541 encoding a globin-like protein for qPCR | 88 bp |
| oCBM4797 | AGCACTGCTGAAGACAATAG | oCBM4798 | AGTTGCCAACCCATCATAAG | Amplification of a fragment of the Mimosa pudica MpudA1P6v1r1_Scf29g0605241 encoding a globin-like protein for qPCR | 94 bp |
| oCBM5685 | CACTTTCGAGGACCAACACG | oCBM5686 | GGAAGGATGGTGTGTGAGTGT | Amplification of a fragment of the Mimosa pudica MpudA1P6v1r1_Scf15g0080361 encoding a putative PR10 protein for qPCR | 84 bp |
| oCBM4991 | TGGGTCAATTCTGCTAGATG | oCBM4992 | TTTGGGAATGCTGTCTTCTC | Amplification of a fragment of the Mimosa pudica MpudA1P6v1r1_Scf04g0255631 ecoding a putative peroxidase for qPCR | 60 bp |
| oCBM5683 | AGCCACCACGATGTTACTGG | oCBM5684 | CCCCAAGATTTGCATGCGTG | Amplification of a fragment of the Mimosa pudica MpudA1P6v1r1_Scf32g0613491 encoding a putative gibberellin 3-beta-dioxygenase for qPCR | 98 bp |
| oCBM4746 | TTGATCTTTGCAGGGAAACAG | oCBM4747 | CGAAGCACAAGATGAAGGGTA | Amplification of a fragment of the Mimosa pudica MpudA1P6v1r1_Scf06g0302791 encoding an ubiquitin-like protein for qPCR normalization | 89 bp |
| oCBM4949 | AAACCCGTATTCAGTCATGGTC | oCBM4950 | AATCCCTCAATGCTCCTTACAC | Amplification of a fragment of the Mimosa pudica MpudA1P6v1r1_Scf39g0686881 encoding a putative helicase for qPCR normalization | 65 bp |
| oCBM4901 | TGCGATGGAAGGGAAAGTAG | oCBM4902 | TCCTCTCACGACACAATCTG | Amplification of a fragment of the Mimosa pudica MpudA1P6v1r1_Scf10g0050861 encoding a putative helicase for qPCR normalization | 89 bp |

### Supplemental materials and methods

#### Genome sequence analysis of evolved clones

Single colonies from purified evolved clones were grown overnight in rich medium supplemented with trimethoprim. Bacterial DNA was extracted from 1 mL of culture using the Wizard genomic DNA purification kit (Promega). Evolved clone DNAs were sequenced at the GeT-PlaGe core facility (https://get. genotoul.fr/), INRAE Toulouse. DNA-seq libraries have been prepared according to Illumina's protocols using the Illumina TruSeq Nano DNA HT Library Prep Kit. Briefly, DNA was fragmented by sonication, size selection was performed using SPB beads (kit beads), and adapters were ligated to be sequenced. Library quality was assessed using an Advanced Analytical Fragment Analyzer (Agilent), and libraries were quantified by quantitative PCR using the Kapa Library Quantification Kit (Roche). Sequencing has been performed on a NovaSeq6000 S4 lane (Illumina) using a paired-end read length of 2 × 150 bp with the Illumina NovaSeq Reagent Kits. Sequencing reads from NovaSeq6000 runs (all whole populations and clones from cycle 35) were mapped on the chimeric reference genome of the ancestral strain, comprising *R. pseudosolanacearum* GMI1000 chromosome (GenBank accession number: NC_003295.1) and megaplasmid (NC_003296.1) together with *C. taiwanensis* symbiotic plasmid pRalta (CU633751). Mutations were detected using breseq v0.33.1 (Deatherage and Barrick 2014) with default parameters using the consensus mode. Mutation lists were curated manually in order to remove mutations present in the ancestral strains as well as false-positive hits arising from reads misalignments in low complexity regions. Mutations detected in evolved clones represented in Fig. 2 are listed in Table S2.

#### Phylogeny of evolved clones

Based on the mutations that occurred during the evolution experiment, artificial sequences were constructed using only the positions mutated in the experiment and assigning each clone to the wild-type or the mutant allele based on its genome sequence. Phylogenetic trees were reconstructed online (www.phylogeny.fr) using the maximum likelihood (ML) heuristic search under the LG model (1) with C-rate variation among sites (2), as implemented in PhyMLv3.0 (3).

#### Mutant construction

The *paeA*$^{V321G}$ SNP mutants were constructed using the MuGent method described previously (4, 5). Briefly, a 6kb fragment carrying the point mutation was amplified by PCR using the G5 evolved clone as DNA matrix and the Phusion high-fidelity DNA polymerase (New England Biolabs). This fragment was co-transformed into competent strains of *R. pseudosolanacearum* strains together with a DNA fragment carrying an antibiotic resistance gene, either spectinomycin (pCBM142) or kanamycin resistance gene (pRCK-*PpsbA*-mCherry or pRCK-*PpsbA*-GFP), integrated into the neutral intergenic region downstream the *glmS* gene. Antibiotic-resistant transformants were selected and screened for the presence of the point mutation using PCR primer pairs specifically amplifying the wild-type or the mutant allele. The mutants were finally verified by Sanger sequencing of the mutated region.

Unmarked deletions of *paeA* and *nifH* were generated using the pEX18Tc plasmid carrying the *sacB* selection gene and the tetracycline resistance gene. Upstream and downstream fragments of the genes to be deleted were amplified by PCR using primers with overhanging *Bam*HI/*Xba*I and XbaI/HindIII restriction sites, digested with the appropriate restriction enzymes and cloned into the pEX18Tc *Bam*HI/*Hin*dIII digested plasmid. The resulting plasmid was introduced into strains of *R. pseudosolanacearum* by natural transformation and selection on tetracycline. The first integration

event was verified by PCR. Transformants were then plated on rich medium supplemented with 5% saccharose to select for the second recombination event. Tetracycline sensitive and saccharose resistant clones were screened by PCR to identify the deleted mutants.
To complement the *paeA* mutants, the wild-type *paeA* gene was amplified by PCR using primers overhanging with *Bam*HI and *Hin*dIII restriction sites and a Phusion high-fidelity DNA polymerase (New England Biolabs). The PCR product was digested with the *Bam*HI and *Hin*dIII restriction enzymes and cloned into the pEX18Tc plasmid digested with the same enzymes. The resulting plasmid was integrated into the native *paeA* locus by natural transformation of *R. pseudosolanacearum* strains and subsequently excised by selection on rich medium supplemented with 5% saccharose. Complemented strains were identified by PCR screening and verified by Sanger sequencing.
To label *R. pseudosolanacearum* strains with a constitutively expressed *lacZ* gene, the P*psbA*-*lacZ* fusion from the pRCG-*PpsbA*-*lacZ* plasmid was integrated in the intergenic region downstream the *glmS* gene by natural transformation.
Plasmids and PCR primers used for strain constructions are listed in Tables S2 and S4.

**Plant dry weights**

The aerial part of plants was harvested at 28 days post inoculation and dried at 65°C for two days. Three independent experiments were performed with 10 measurements per experiment and inoculated strain.

***M. pudica* RNA extraction and gene expression analysis**

Nodules were harvested at 10 days post-inoculation and ground in liquid nitrogen using a mortar and pestle prior to RNA extraction. To optimize the grinding, the powder was further mixed in a bead mill (Retsch MM400) for 30 sec 2 times at 30 Hz. Total plant RNA was isolated using the NucleoSpin RNA Plus kit (Macherey-Nagel) according to the manufacturer's instructions, treated with a DNAse (Invitrogen) for 30 min at 37°C and then cleaned up with the NucleoSpin RNA clean-up kit (Macherey-Nagel). RNA quality was verified using a 2100 Bioanalyzer instrument (Agilent) and quantified using a QubitTM fluorometer (Thermo Fisher Scientific). cDNAs were synthesized from 1 µg of extracted RNA using a Transcriptor Reverse Transcriptase kit (Roche, Life technologies) and random hexamers as primers. Finally, *M. pudica* gene expression was measured using the Takyon® No ROX SYBR 2X MasterMix blue dTTP (Eurogentec) and the CFX Opus Dx Real-Time PCR Detection Systems (Bio-rad). Gene expression was normalized by the expression of three housekeeping genes encoding a putative ubiquitin protein (MpudA1P6v1r1_Scf06g0302791) and two putative DNA helicases (MpudA1P6v1r1_Scf39g0686881 and MpudA1P6v1r1_Scf10g0050861).

**Supplemental references**


1. Le SQ, Gascuel O. 2008. An improved general amino acid replacement matrix. *Mol Biol Evol* 25:1307-20.
2. Yang Z. 1994. Maximum likelihood phylogenetic estimation from DNA sequences with variable rates over sites: approximate methods. *J Mol Evol* 39:306-14.
3. Guindon S, Dufayard JF, Lefort V, Anisimova M, Hordijk W, Gascuel O. 2010. New algorithms and methods to estimate maximum-likelihood phylogenies: assessing the performance of PhyML 3.0. *Syst Biol* 59:307-21.

4. Dalia AB, McDonough E, Camilli A. 2014. Multiplex genome editing by natural transformation. *Proc Natl Acad Sci U S A* 111:8937-8942.
5. Capela D, Marchetti M, Clérissi C, Perrier A, Guetta D, Gris C, Valls M, Jauneau A, Cruveiller S, Rocha EPC, Masson-Boivin C. 2017. Recruitment of a lineage-specific virulence regulatory Pathway promotes intracellular infection by a plant pathogen experimentally evolved into a legume symbiont. *Mol Biol Evol* 34:2503-2521.
